## Supplementary figures for "Dimeric G-quadruplex motifs-induced NFRs determine strong replication origins in vertebrates"

A

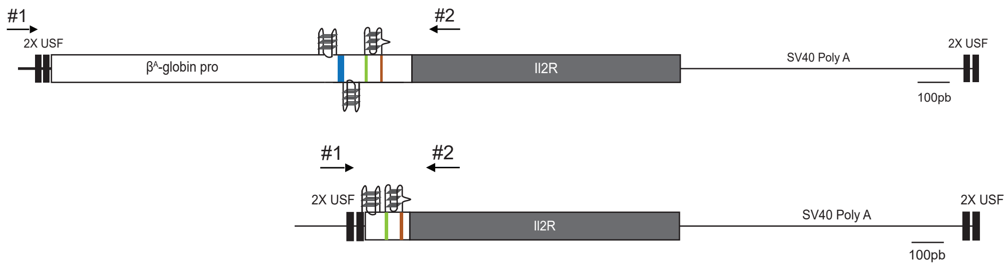


B

| Relative quantification values | | *Med14* gene | *IL2R* gene |
|---|---|---|---|
| $RT^+$-qPCR | $\beta^A$-globin full origin (1) | 100 | 0.04 |
| | $\beta^A$-globin full origin (2) | 100 | 0.03 |
| | $\beta^A$-globin minimal origin (1) | 100 | 0.04 |
| | $\beta^A$-globin minimal origin (2) | 100 | 0.01 |

| Crossing point values | | *Med14 gene* | *IL2R gene* |
|---|---|---|---|
| $RT^+$-qPCR | $\beta^A$-globin full origin (1) | 22.28 | 35.41 |
| | $\beta^A$-globin full origin (2) | 21.38 | 35.14 |
| $RT^-$-qPCR | $\beta^A$-globin full origin (1) | NA | NA |
| | $\beta^A$-globin full origin (2) | NA | NA |
| $RT^+$-qPCR | $\beta^A$-globin minimal origin (1) | 22.03 | 35.27 |
| | $\beta^A$-globin minimal origin (2) | 21.03 | 36.46 |
| $RT^-$-qPCR | $\beta^A$-globin minimal origin (1) | NA | NS |
| | $\beta^A$-globin minimal origin (2) | NA | NA |

C

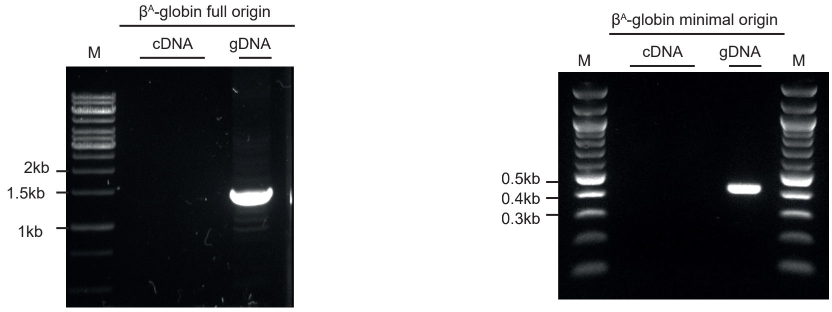


**Supplementary Figure 1: The full and minimal $\beta^A$-globin origins do not drive transcription**

(A) Schemes of the full and minimal βA-globin origins coupled with the IL2R gene, SV40 PolyA sequence and USF binding sites are shown. Positions of specific primers (#1 and #2) used to detect potential RNAs elongated over pG4s #1 and #3 are indicated with arrows. (B) Relative quantification by RT-qPCR on mRNAs was performed in clonal cell lines containing the full and minimal $\beta^A$-globin origins. mRNA levels were normalized against Med14 mRNA levels arbitrarily set at 100 (first table). Crossing point values for $RT^+$-qPCR and $RT^-$-qPCR (background levels) experiments are reported in the second table. NA and NS correspond to non-amplified and non-specific signals respectively. (C) PCR products obtained with primers #1 and #2 to amplify either cDNA (cDNA) or genomic DNA (gDNA) as a control were subjected to electrophoresis in a 1% w/v agarose gel and stained with SYBR safe. The DNA size marker used is a commercial 1 kb plus DNA ladder (M, left panel) or a 100 bp DNA ladder (M, right panel). The absence of the 1.3 kb or the 0.4 kb PCR products after amplification over pG4 #1 and #3 with primers #1 and #2 in cDNAs from the full and minimal $\beta^A$-globin origins demonstrates the absence of transcription at these sites.

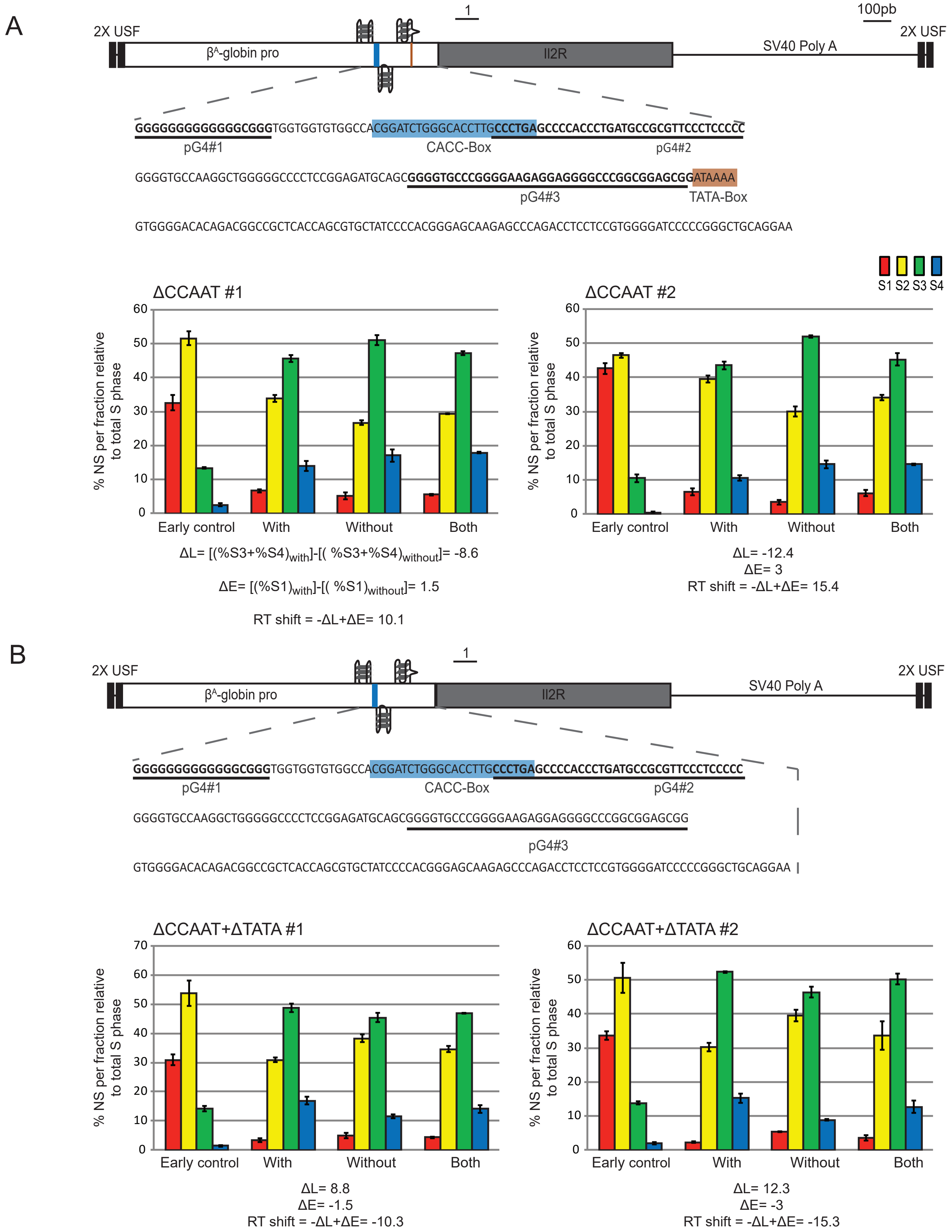


**Supplementary Figure 2: RT analyses of the ΔCCAAT and ΔCCAAT+ΔTATA β$^{A}$-globin origins**

(A-B) Schemes of the ΔCCAAT β$^{A}$-globin origin (A) and the ΔCCAAT+ΔTATA βA-globin origin (B) coupled with the IL2R gene, SV40 PolyA sequence and USF binding sites are shown on the top. The corresponding sequences are given below each scheme with pG4s underlined and cis-regulatory elements delineated by colored boxes (blue for the CACC-Box and brown for the TATA-Box). Analyses of two ΔCCAAT βA-globin origin clonal cell lines (A) or two ΔCCAAT+ΔTATA β$^{A}$-globin origin clonal cell lines (B) are reported. RT profiles of each chromosomal allele are determined after targeted transgene integration using the allele-specific analysis of RT method by real-time PCR quantification (1). BrdU pulse-labeled cells were sorted into four S-phase fractions from early to late (S1 to S4) and the immune-precipitated newly synthesized strands (NS) were quantified by real-time qPCR in each fraction. Specific primer pairs determine the RT profile for the modified allele (With, black line, 1), the wt allele (Without) and both alleles (Both). The endogenous β-globin locus was analyzed as an early-replicated control (Early). Differences in -ΔL+ΔE values calculated at the target site following transgene integration are indicated. Error bars correspond to the standard deviation for qPCR duplicates.

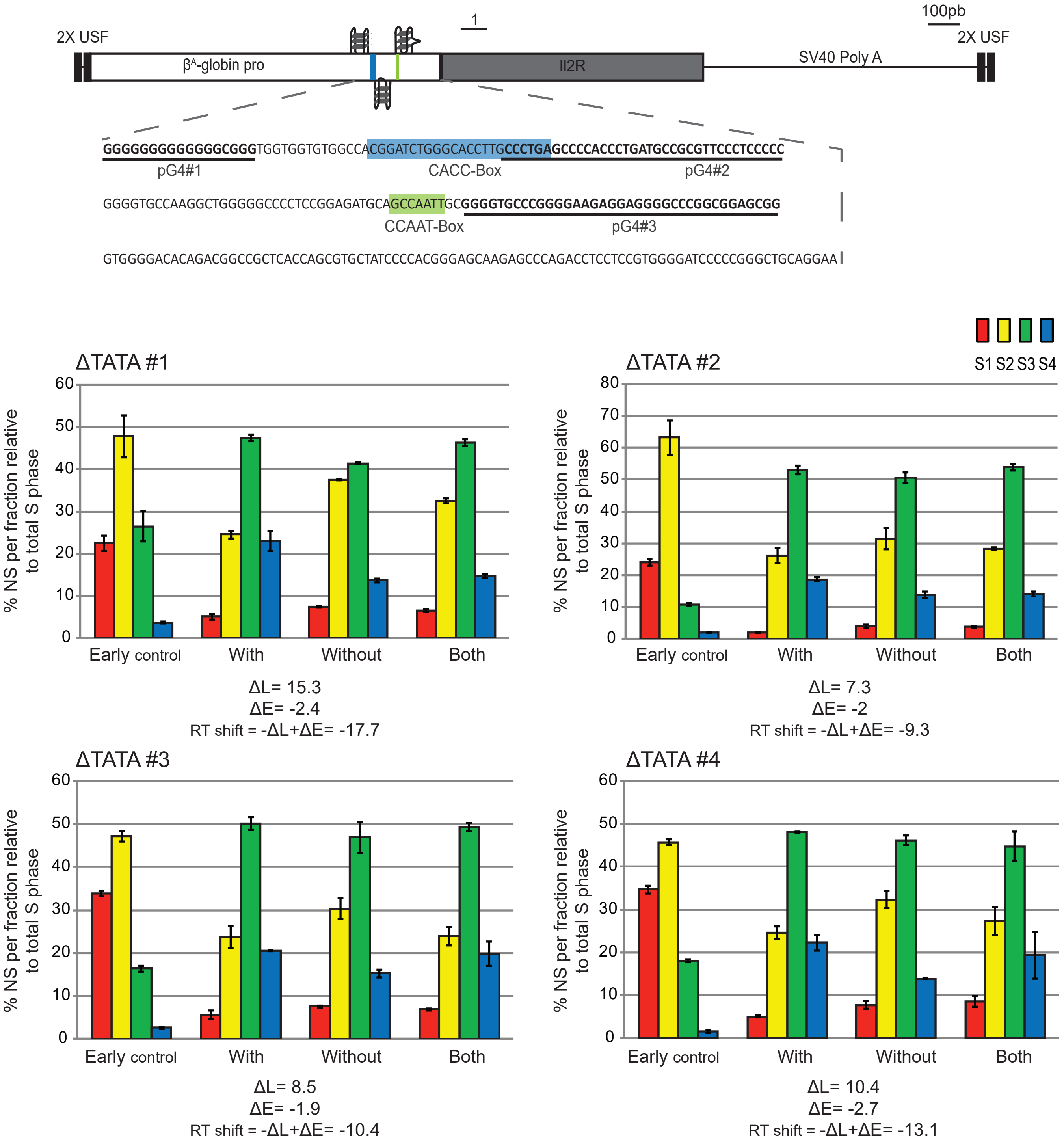


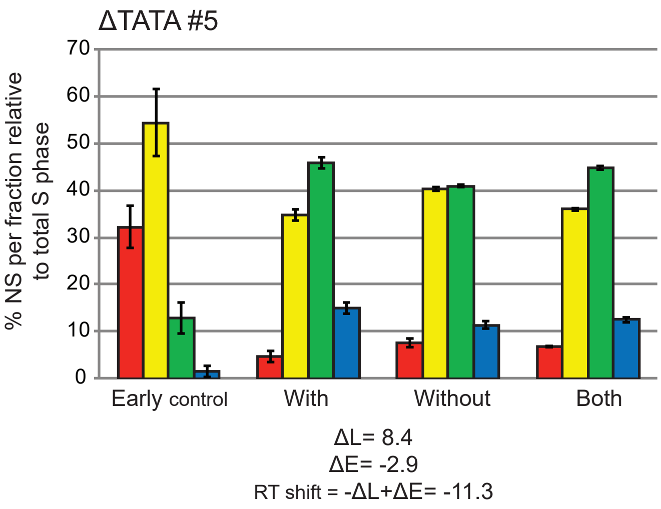


**Supplementary Figure 3: RT analyses of the ΔTATA β$^{A}$-globin origin**
Scheme of the β$^{A}$-globin ΔTATA origin coupled with the IL2R gene, SV40 PolyA sequence and USF binding sites is shown on the top. The corresponding sequence is given below with pG4s underlined and cis-regulatory elements delineated by colored boxes (blue for the CACC-Box and green for the CCAAT-Box). Analyses of five ΔTATA β$^{A}$-globin origin clonal cell lines are reported. RT profiles of each chromosomal allele is determined after targeted transgene integration using the allele-specific analysis of RT method by real-time PCR quantification as described in supplementary figure 2.

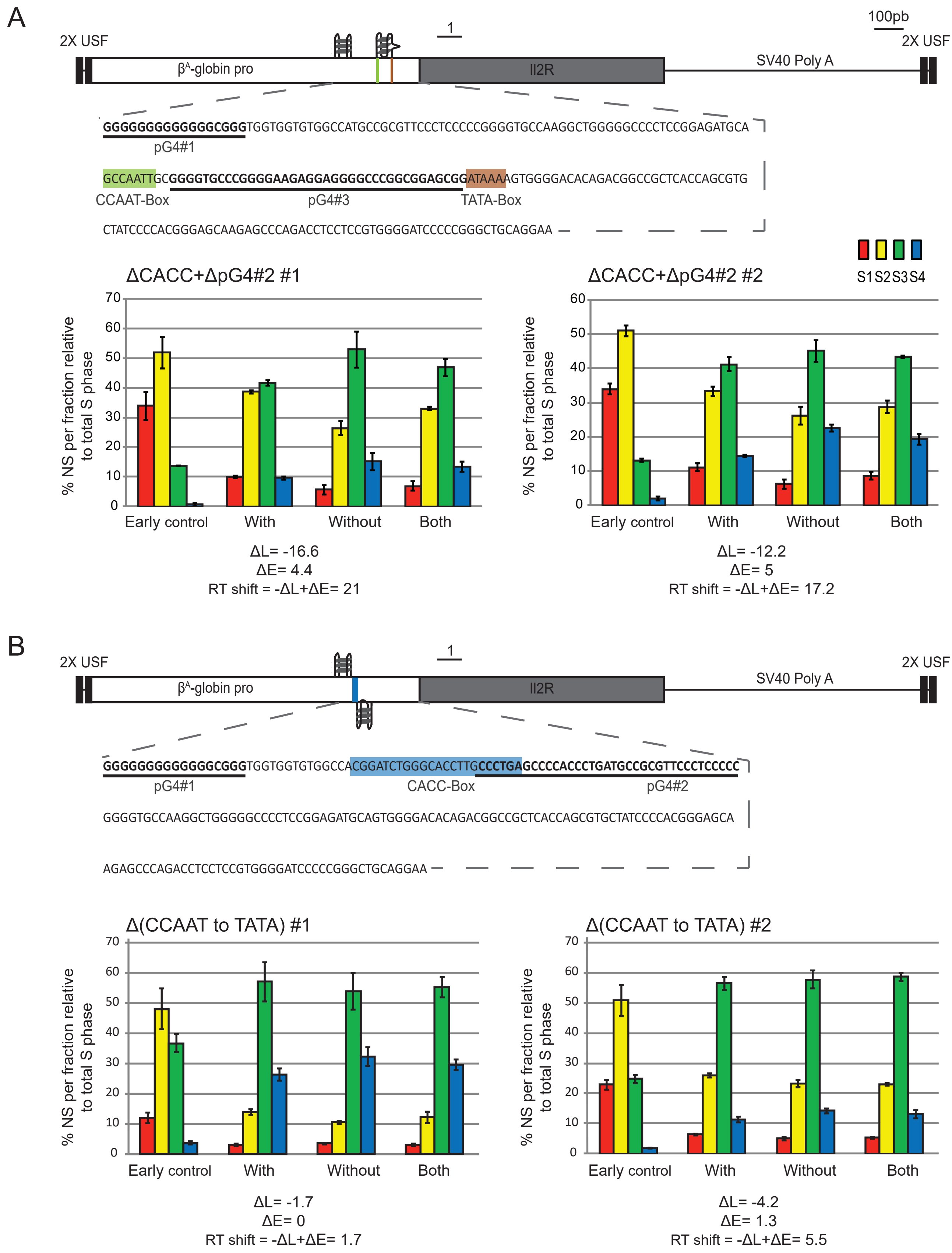


**Supplementary Figure 4: RT analyses of the ΔCACC+pG4#2 and Δ(CCAAT to TATA) $\beta^A$-globin origins**
(A-B) Schemes of the $\beta^A$-globin ΔCACC+pG4#2 origin (A) and the $\beta^A$-globin Δ(CCAAT to TATA) origin (B) coupled with the IL2R gene, SV40 PolyA sequence and USF binding sites are shown on the top. The corresponding sequences are given below each scheme with pG4s underlined and cis-regulatory elements delineated by colored boxes (blue for the CACC-Box, green for the CCAAT-Box and brown for the TATA-Box). Analyses of two ΔCAACC+pG4#2 $\beta^A$-globin origin clonal cell lines (A) or two Δ(CCAAT to TATA) $\beta^A$-globin origin clonal cell lines (B) are reported. RT profiles of each chromosomal allele are determined after targeted transgene integration using the allele-specific analysis of RT method by real-time PCR quantification as described in supplementary figure 2.

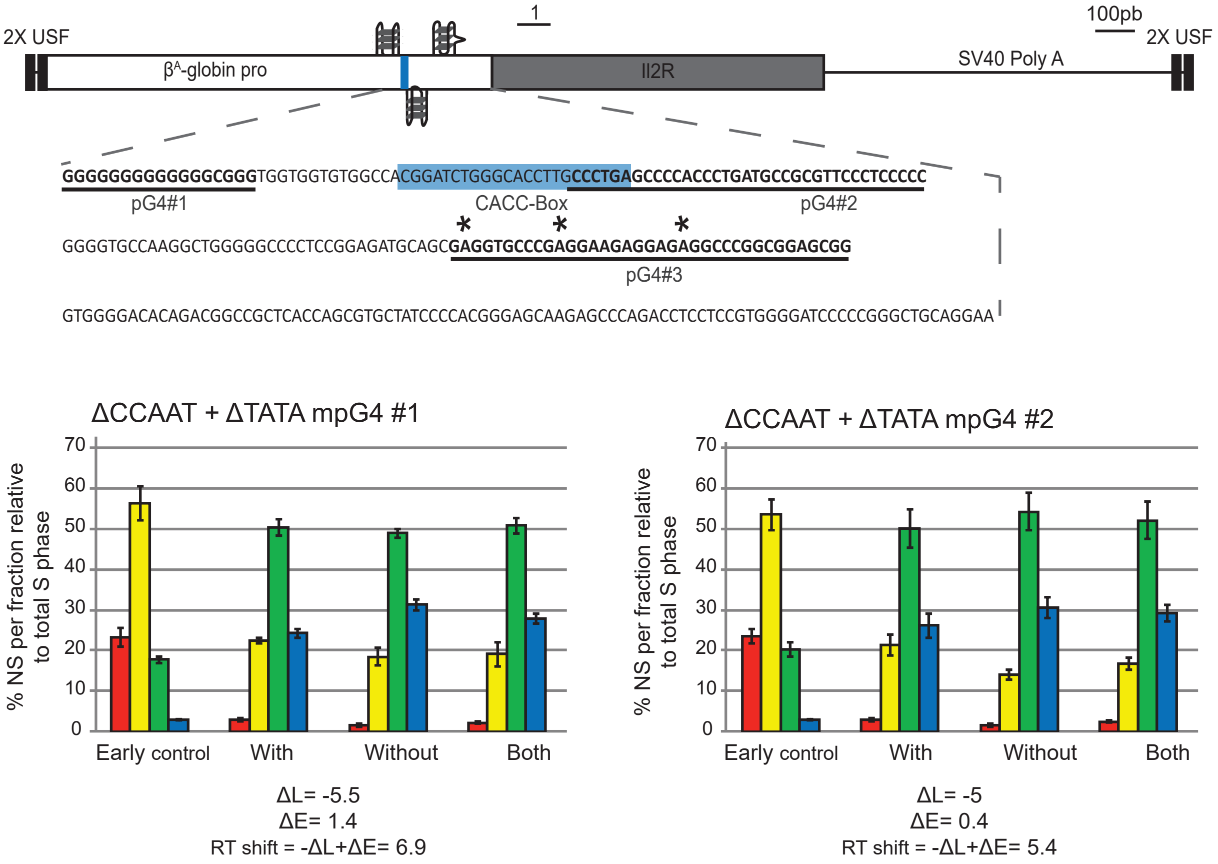


**Supplementary Figure 5: RT analyses of the ΔCCAAT+ΔTATA mpG4 $\beta^{A}$-globin origin**

Scheme of the $\beta^{A}$-globin ΔCCAAT+ΔTATA mpG4 origin coupled with the IL2R gene, SV40 PolyA sequence and USF binding sites is shown at the top. The corresponding sequence is given below with pG4s underlined and cis-regulatory elements delineated by colored boxes (blue for the CACC-Box). Points mutation inside the pG4#3 are indicated with black stars. Analyses of two ΔCCAAT+ΔTATA $\beta^{A}$-globin origin clonal cell lines are reported. RT profiles of each chromosomal allele is determined after targeted transgene integration using the allele-specific analysis of RT method by real-time PCR quantification as described in supplementary figure 2.

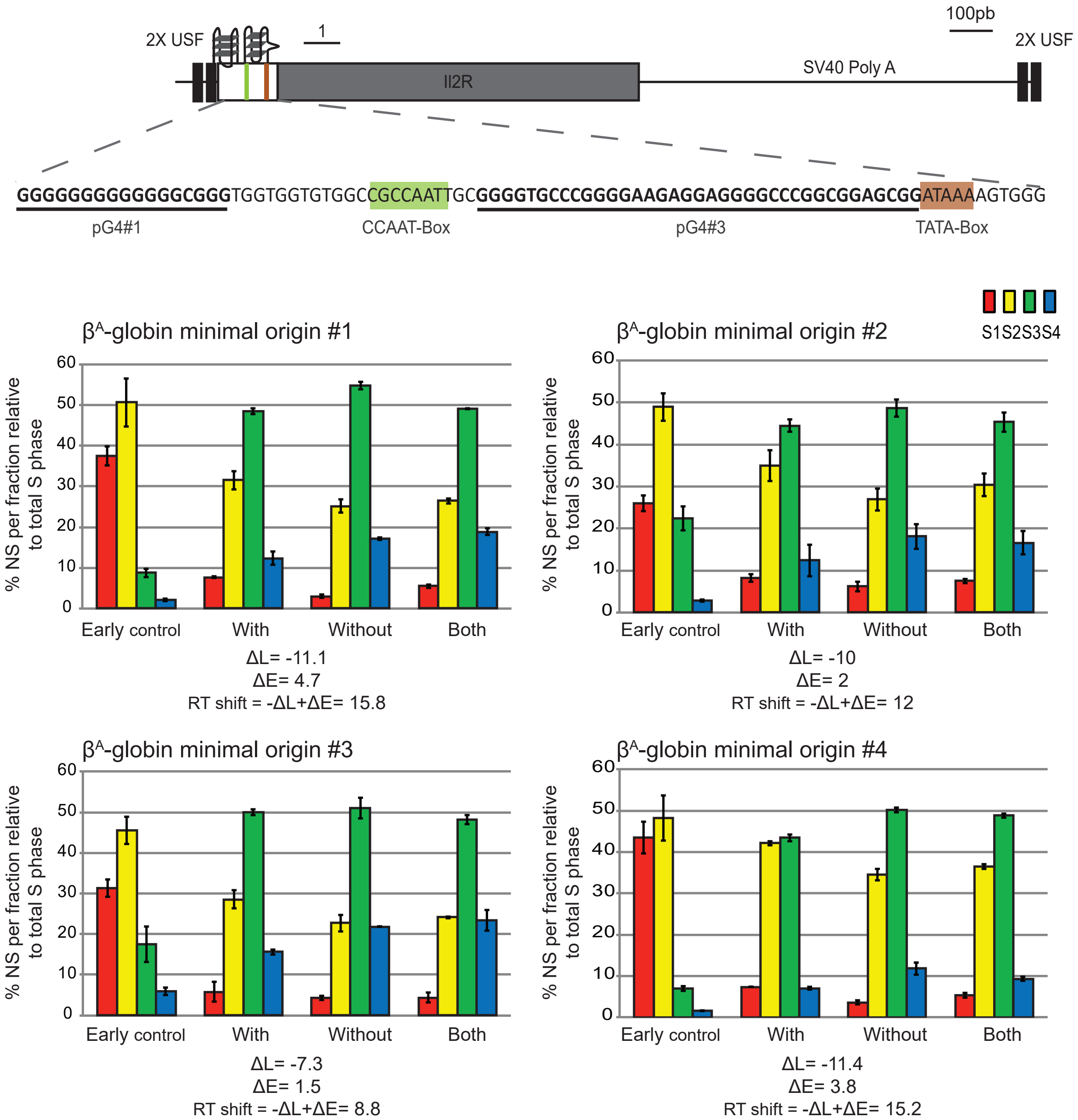


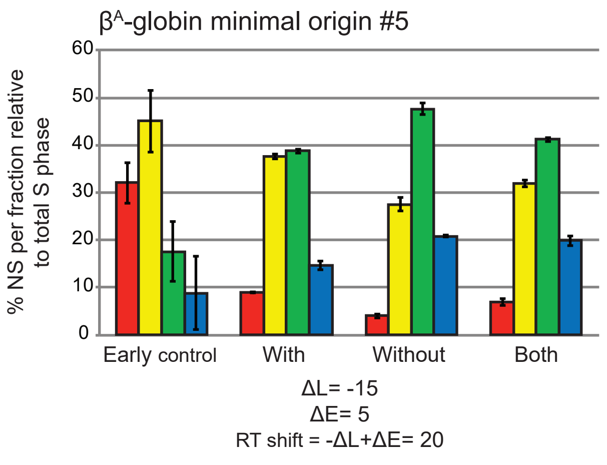


**Supplementary Figure 6: RT analyses of the $\beta^A$-globin minimal origin**

Scheme of the $\beta^A$-globin minimal origin coupled with the IL2R gene, SV40 PolyA sequence and USF binding sites is shown at the top. The corresponding sequence is given below each scheme with pG4s underlined and cis-regulatory elements delineated by colored boxes (green for the CCAAT-Box and brown for TATA-Box). Analyses of five $\beta^A$-globin minimal origin clonal cell lines are reported. RT profiles of each chromosomal allele is determined after targeted transgene integration using the allele-specific analysis of RT method by real-time PCR quantification as described in supplementary figure 2.

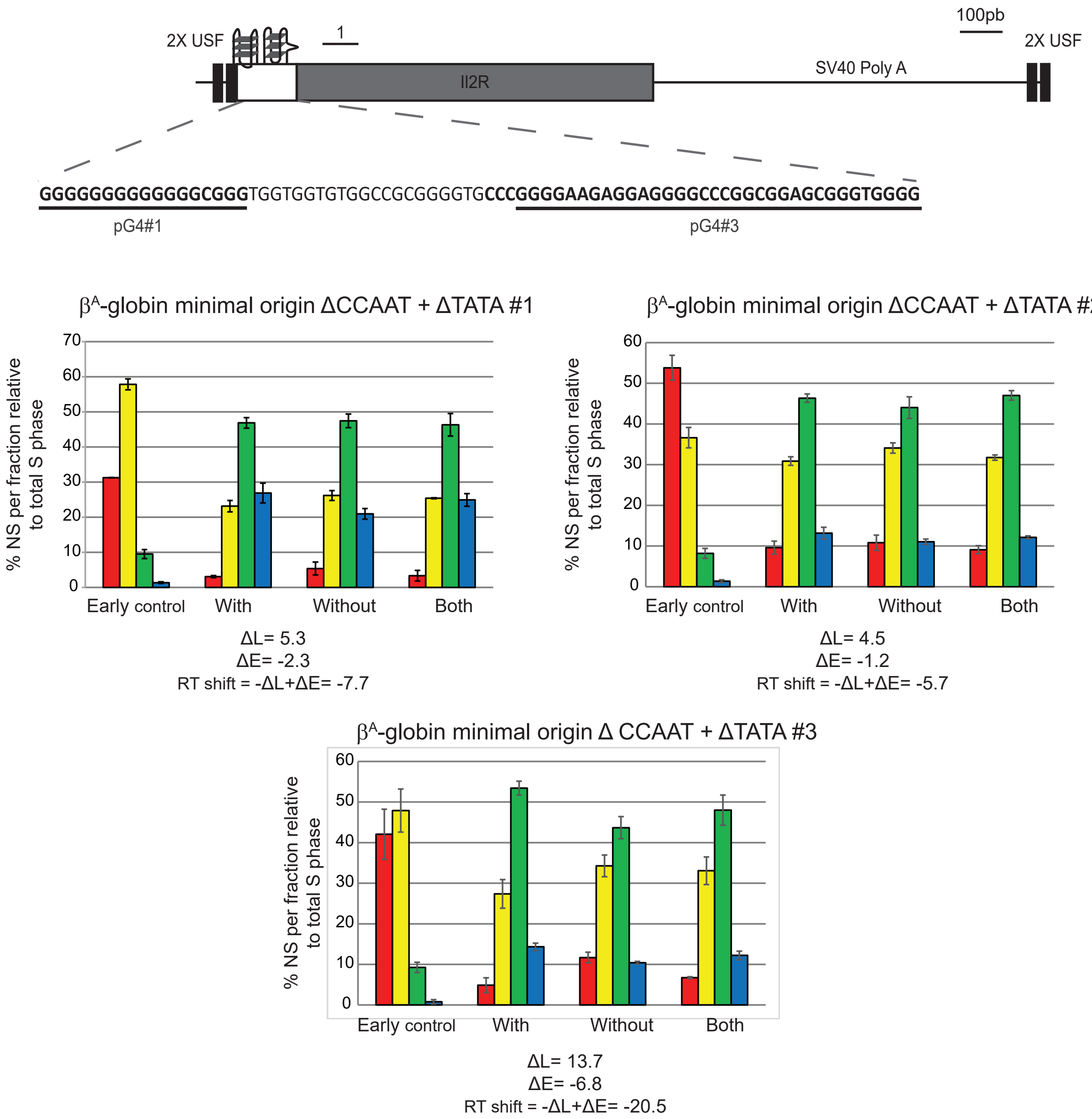


**Supplementary Figure 7: RT analyses of the $\beta^A$-globin minimal origin ΔCCAAT+ΔTATA**
Scheme of the $\beta^A$-globin minimal origin ΔCCAAT+ΔTATA coupled with the IL2R gene, SV40 PolyA sequence and USF binding sites is shown at the top. The corresponding sequence is given below with pG4s underlined. Analyses of three $\beta^A$-globin minimal origin ΔCCAAT+ΔTATA clonal cell lines are reported. RT profiles of each chromosomal allele is determined after targeted transgene integration using the allele-specific analysis of RT method by real-time PCR quantification as described in supplementary figure 2.

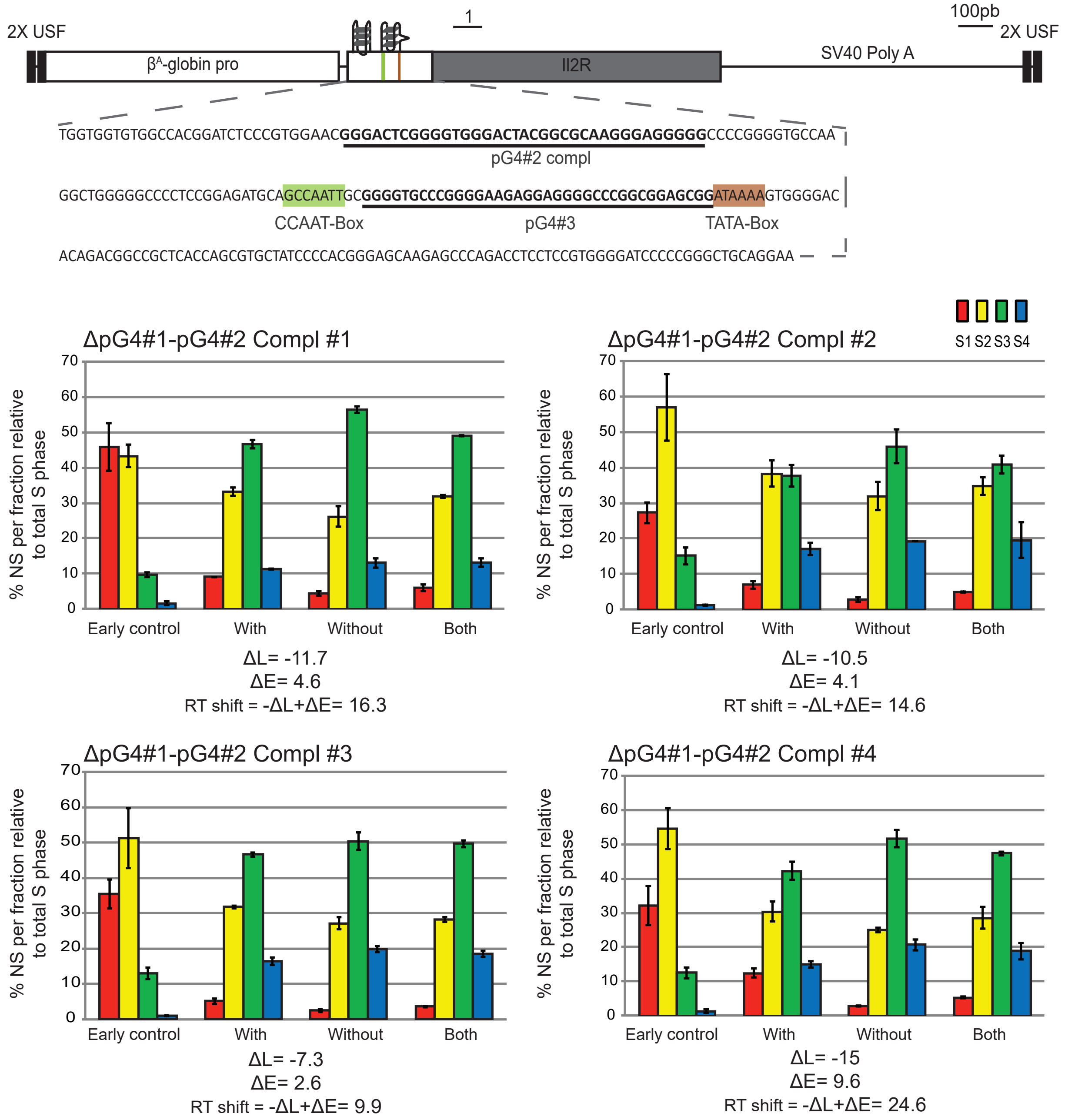


**Supplementary Figure 8: RT analyses of the ΔpG4#1 pG4#2 Compl $\beta^{A}$-globin origin**

Scheme of the ΔpG4#1 pG4#2 Compl $\beta^{A}$-globin origin coupled with the IL2R gene, SV40 PolyA sequence and USF binding sites is shown at the top. The corresponding sequence is given below with pG4s underlined and cis-regulatory elements delineated by colored boxes (green for the CCAAT-Box and brown for TATA-Box). Analyses of four ΔpG4#1 pG4#2 Compl $\beta^{A}$-globin origin clonal cell lines are reported. RT profiles of each chromosomal allele is determined after targeted transgene integration using the allele-specific analysis of RT method by real-time PCR quantification as described in supplementary figure 2.

A

Synchronized $\beta^A$-globin minimal origin

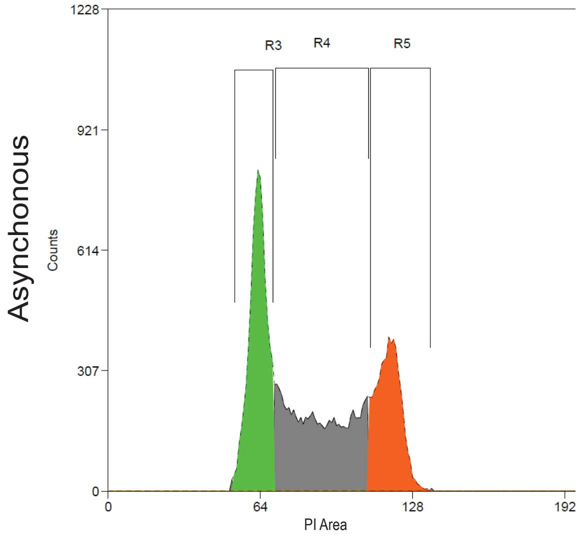


| Region | Count | % Hist | Median | Mean | CV |
|---|---|---|---|---|---|
| Total | 20000 | 100.00 | 83.00 | 86.83 | 26.34 |
| R3 | 7151 | 35.75 | 63.00 | 62.82 | 5.62 |
| R4 | 7789 | 38.95 | 88.00 | 88.80 | 13.68 |
| R5 | 5024 | 25.12 | 118.00 | 118.07 | 4.03 |

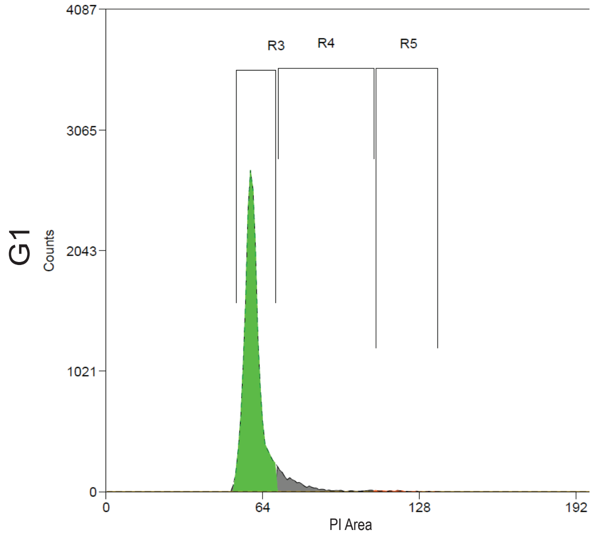


| Region | Count | % Hist | Median | Mean | CV |
|---|---|---|---|---|---|
| Total | 20000 | 100.00 | 60.00 | 61.83 | 12.92 |
| R3 | 18009 | 90.05 | 60.00 | 59.84 | 5.29 |
| R4 | 1801 | 9.01 | 75.00 | 77.97 | 10.95 |
| R5 | 128 | 0.64 | 118.00 | 118.34 | 5.27 |

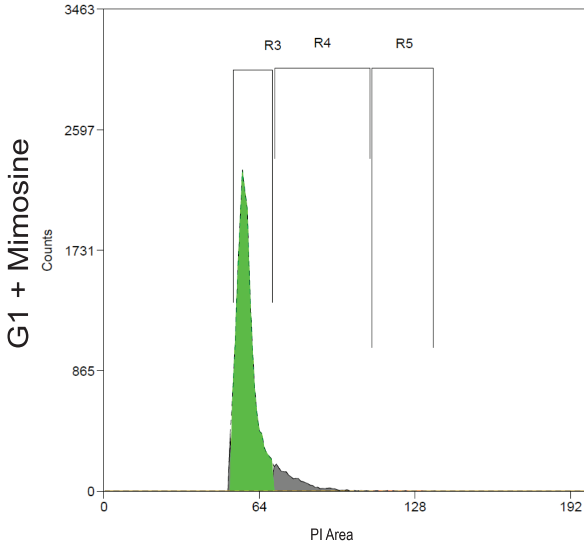


| Region | Count | % Hist | Median | Mean | CV |
|---|---|---|---|---|---|
| Total | 20000 | 100.00 | 58.00 | 60.55 | 12.62 |
| R3 | 17523 | 87.61 | 58.00 | 58.61 | 6.13 |
| R4 | 2033 | 10.17 | 76.00 | 77.99 | 9.37 |
| R5 | 31 | 0.16 | 117.00 | 119.03 | 4.99 |

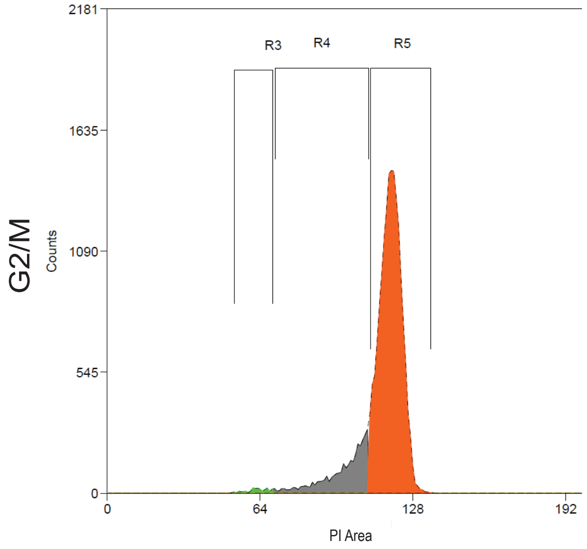


| Region | Count | % Hist | Median | Mean | CV |
|---|---|---|---|---|---|
| Total | 20000 | 100.00 | 118.00 | 114.84 | 9.33 |
| R3 | 222 | 1.11 | 63.00 | 62.67 | 6.77 |
| R4 | 3292 | 16.46 | 102.00 | 98.92 | 9.53 |
| R5 | 16479 | 82.39 | 119.00 | 118.73 | 3.59 |

**Supplementary Figure 9: Validation of the $\beta^A$-globin minimal origin cell synchronization by elutriation and L-mimosine incubation**

Cell cycle analyses by flow cytometry for different fractions of elutriated cells containing the $\beta^A$-globin minimal origin were made after DNA labelling with propidium iodide (PI). The DNA content distributions of cells in G1-phase before (G1) and G1 plus 3 hrs of L-mimosine incubation (G1+Mimosine) or in G2/M-phase (G2/M) are shown. G1-phase cells are represented in green and G2/M cells in orange. R3, R4 and R5 regions were settled based on the asynchronous cell profile to define the proportion of cells in each fraction. Proportions of cells analyzed for each condition as well as statistics of the distribution of the detected PI values for each region are given in the table below each graph. Black titles on the Y-axis give the name of the cell population analyzed.

B

### Synchronised β$^A$-globin minimal origin pG4#1 rev compl

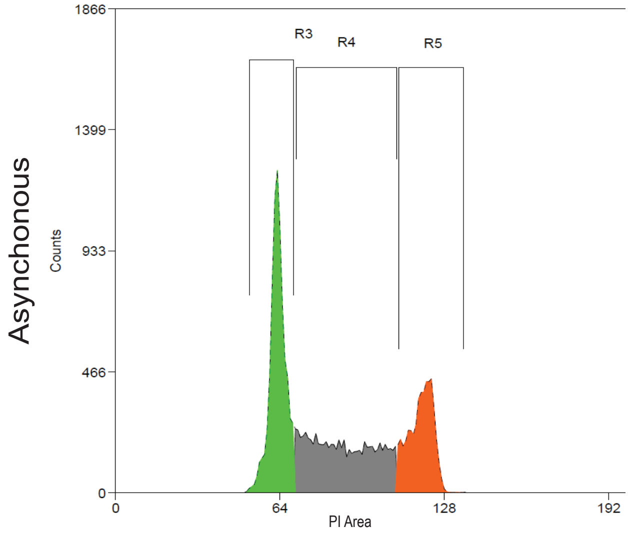


| Region | Count | % Hist | Median | Mean | CV |
|---|---|---|---|---|---|
| Total | 20000 | 100.00 | 80.00 | 86.08 | 26.74 |
| R3 | 7704 | 38.52 | 63.00 | 63.02 | 4.78 |
| R4 | 7393 | 36.96 | 88.00 | 88.43 | 13.36 |
| R5 | 4896 | 24.48 | 119.00 | 118.82 | 3.76 |

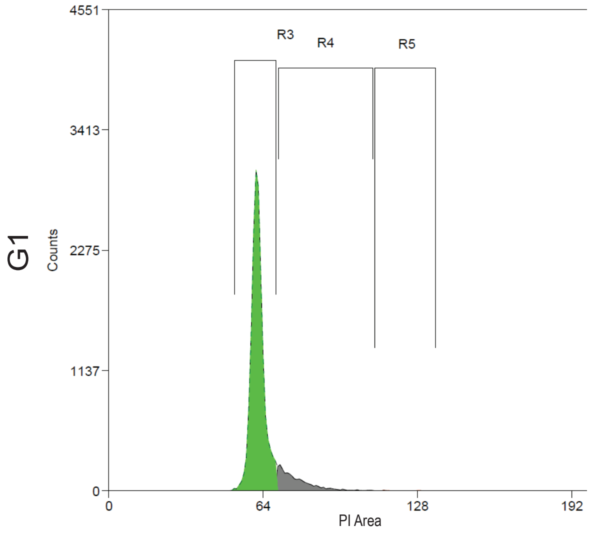


| Region | Count | % Hist | Median | Mean | CV |
|---|---|---|---|---|---|
| Total | 19189 | 100.00 | 62.00 | 63.75 | 10.35 |
| R3 | 16785 | 87.47 | 62.00 | 61.71 | 4.31 |
| R4 | 2379 | 12.40 | 76.00 | 77.65 | 9.31 |
| R5 | 20 | 0.10 | 116.00 | 119.30 | 5.35 |

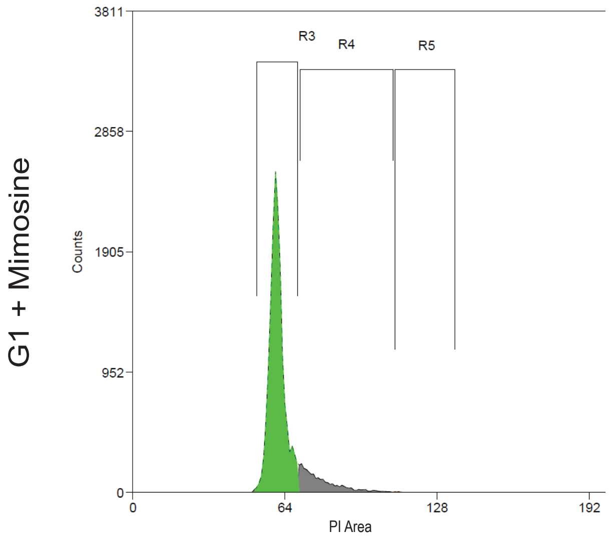


| Region | Count | % Hist | Median | Mean | CV |
|---|---|---|---|---|---|
| Total | 18386 | 100.00 | 61.00 | 63.26 | 12.47 |
| R3 | 15790 | 85.88 | 60.00 | 60.64 | 5.06 |
| R4 | 2554 | 13.89 | 77.00 | 79.13 | 10.62 |
| R5 | 23 | 0.13 | 112.00 | 114.09 | 4.12 |

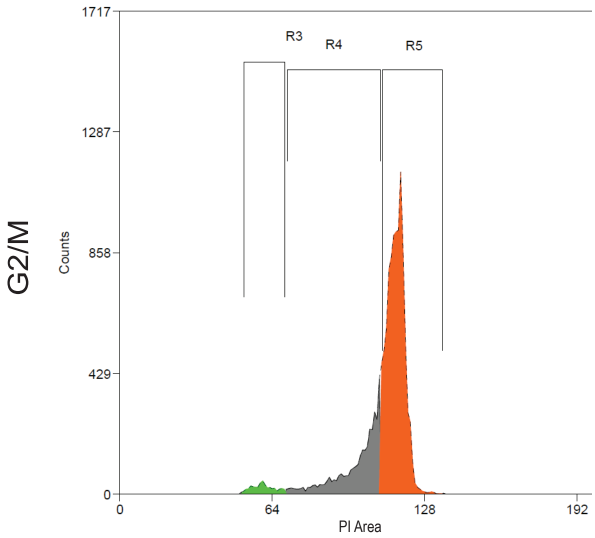


| Region | Count | % Hist | Median | Mean | CV |
|---|---|---|---|---|---|
| Total | 13150 | 100.00 | 114.00 | 109.93 | 11.82 |
| R3 | 422 | 3.21 | 60.00 | 60.27 | 7.58 |
| R4 | 3406 | 25.90 | 103.00 | 99.48 | 9.67 |
| R5 | 9312 | 70.81 | 116.00 | 116.04 | 2.96 |

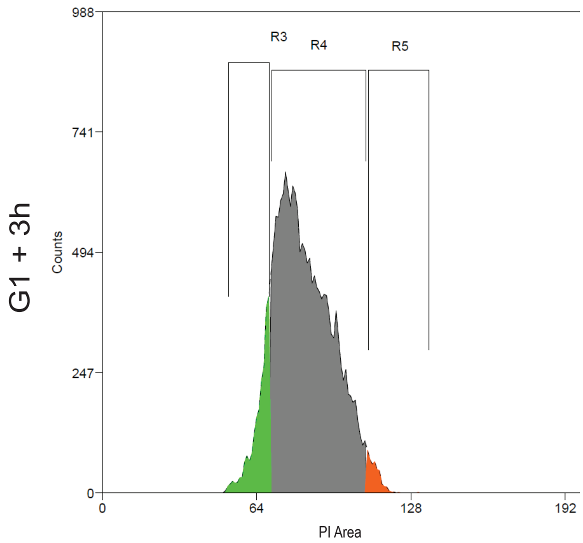


| Region | Count | % Hist | Median | Mean | CV |
|---|---|---|---|---|---|
| Total | 18685 | 100.00 | 82.00 | 83.42 | 14.74 |
| R3 | 2161 | 11.57 | 66.00 | 65.16 | 5.96 |
| R4 | 16093 | 86.13 | 84.00 | 85.10 | 11.90 |
| R5 | 427 | 2.29 | 112.00 | 112.83 | 2.27 |

**Supplementary Figure 10: Validation of cells containing the β$^A$-globin minimal inactive origin synchronization by elutriation and L-mimosine incubation**

Cell cycle analyses by flow cytometry of different fractions of elutriated of cells containing the β$^A$-globin minimal origin pG4#1 reverse complement (β$^A$-globin minimal origin pG4#1 rev compl) were made as described in supplementary figure 9. G1-phase cells released for 3 hrs in normal culture conditions were also tested as control of the cell population viability after elutriation.

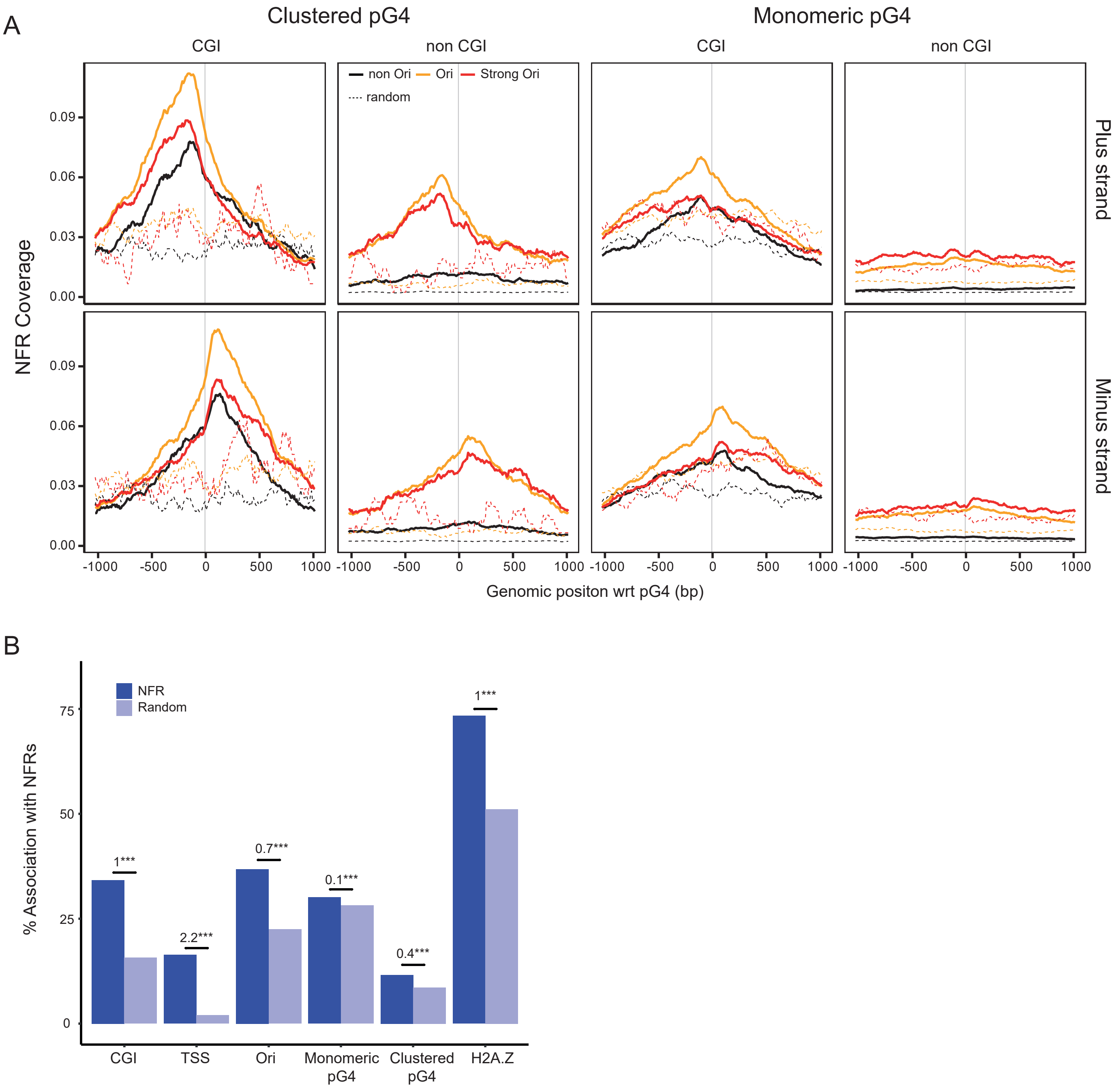


**Supplementary Figure 11**

(A) NFR coverage in G2/M cells around clustered pG4s (left) and monomeric pG4s (right) on the plus (top) and minus (bottom) strand. (B) Association of NFRs with genomic features in chicken cells.

**Supplementary Figure 12**

(A) Nucleosome coverage around cluster (left) and monomeric (right) pG4s on the minus strand. (B) NFR coverage in G1 cells around cluster (left) and monomeric (right) pG4s on the minus strand. (C) Short Nascent Strand coverage around cluster (left) and monomeric (right) pG4s on the minus strand. (D) Cluster (left) and monomeric (right) pG4s coverage around NFR centers on the minus strand. (E) H2A.Z coverage around origins' centers containing cluster (left) or monomeric (right) pG4s found on either the plus (top) or the minus (bottom) strand.

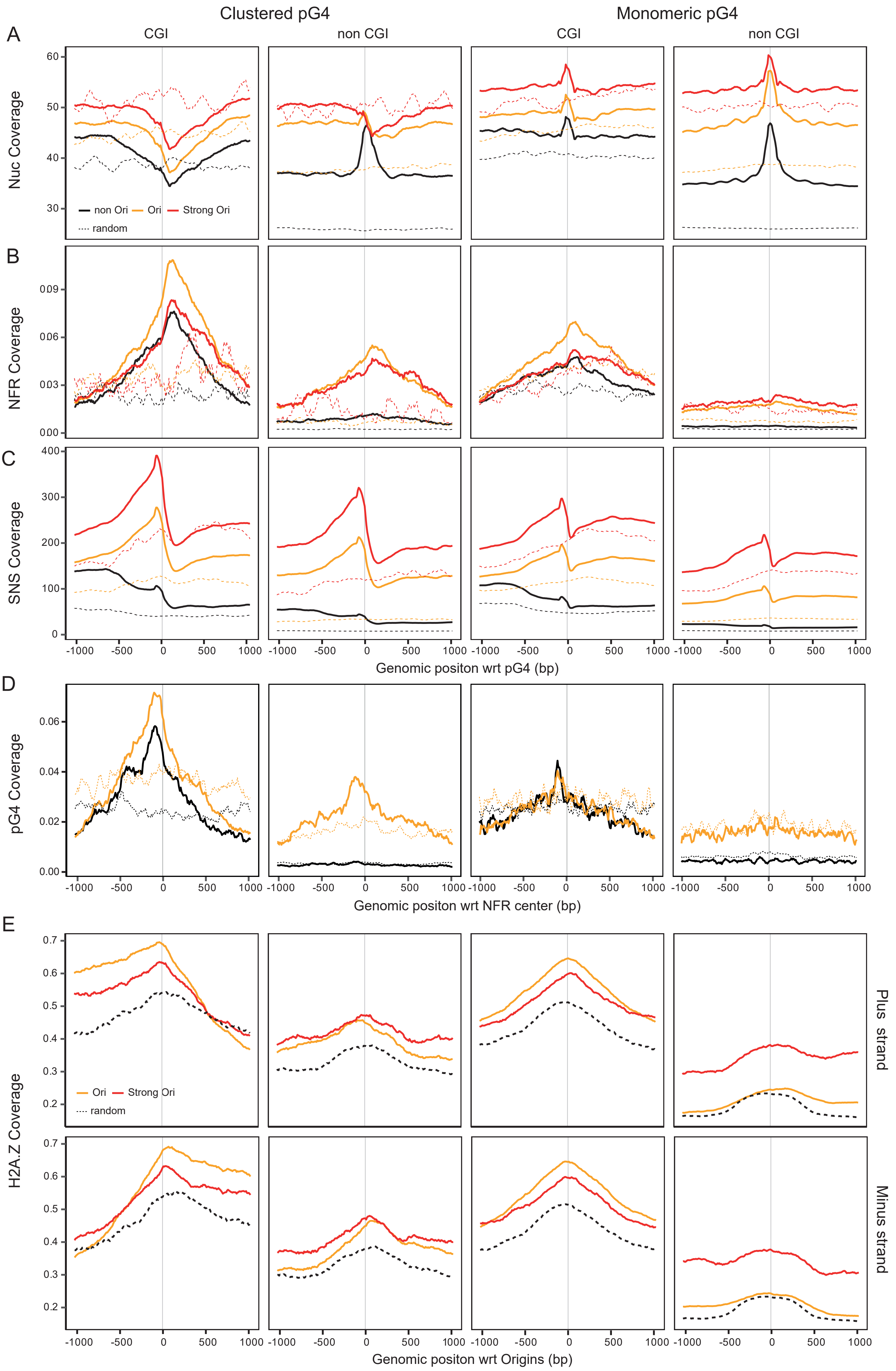

Clustered pG4
Monomeric pG4
A
CGI
non CGI
CGI
non CGI
Nuc Coverage
60
50
40
30
non Ori
Ori
Strong Ori
random
B
NFR Coverage
009
006
003
000
C
SNS Coverage
400
300
200
100
0
-1000
-500
0
500
1000
Genomic positon wrt pG4 (bp)
D
pG4 Coverage
0.06
0.04
0.02
0.00
Genomic positon wrt NFR center (bp)
E
H2A.Z Coverage
0.7
0.6
0.5
0.4
0.3
0.2
Ori
Strong Ori
random
Plus strand
Minus strand
Genomic positon wrt Origins (bp)

### Synchronised wt DT40 cells

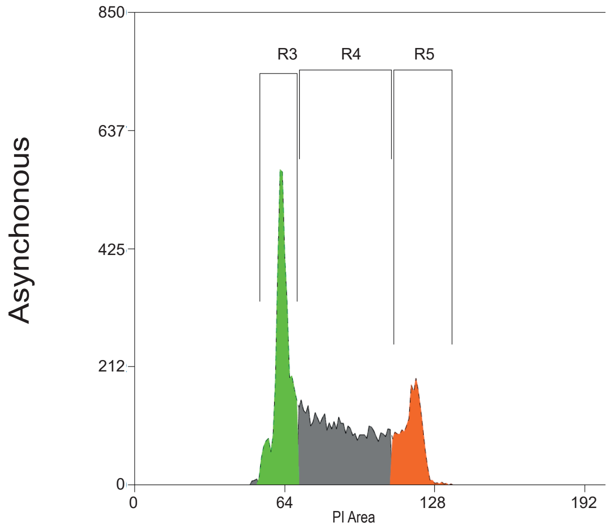


| Region | Count | % Hist | Median | Mean | CV |
|---|---|---|---|---|---|
| Total | 9752 | 100.00 | 79.00 | 84.27 | 25.48 |
| R3 | 3611 | 37.03 | 63.00 | 62.85 | 5.28 |
| R4 | 4258 | 43.66 | 87.00 | 87.97 | 13.31 |
| R5 | 1857 | 19.04 | 118.00 | 117.87 | 3.64 |

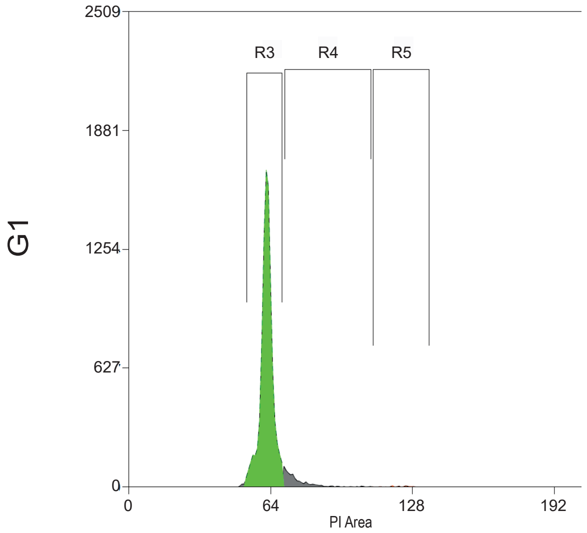


| Region | Count | % Hist | Median | Mean | CV |
|---|---|---|---|---|---|
| Total | 9909 | 100.00 | 62.00 | 63.46 | 10.44 |
| R3 | 9100 | 91.84 | 62.00 | 62.15 | 4.54 |
| R4 | 722 | 7.29 | 74.00 | 76.41 | 9.92 |
| R5 | 50 | 0.50 | 122.00 | 122.08 | 4.65 |

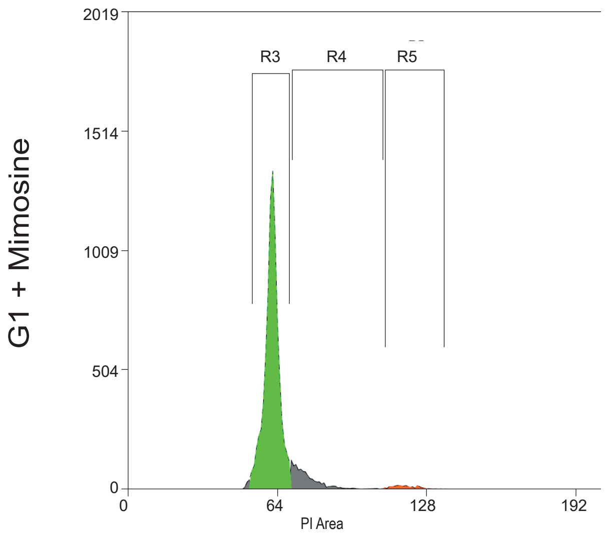


| Region | Count | % Hist | Median | Mean | CV |
|---|---|---|---|---|---|
| Total | 9517 | 100.00 | 62.00 | 64.27 | 15.39 |
| R3 | 8215 | 86.32 | 62.00 | 61.57 | 5.02 |
| R4 | 1033 | 10.85 | 75.00 | 76.61 | 8.62 |
| R5 | 191 | 2.01 | 118.00 | 118.80 | 4.29 |

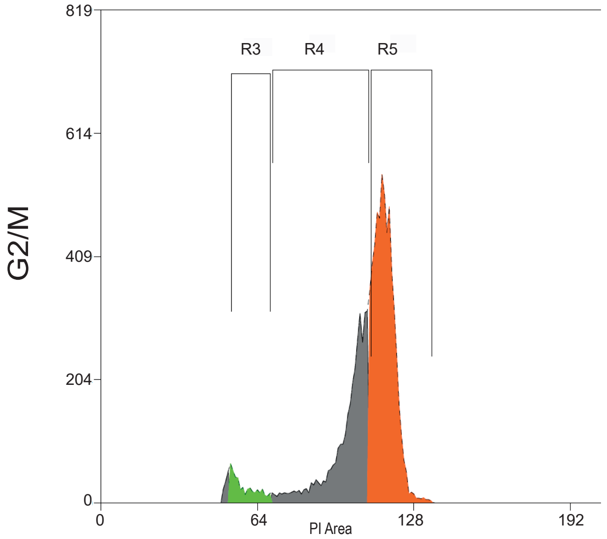


| Region | Count | % Hist | Median | Mean | CV |
|---|---|---|---|---|---|
| Total | 9527 | 100.00 | 112.00 | 107.09 | 14.70 |
| R3 | 452 | 4.74 | 57.00 | 58.79 | 8.41 |
| R4 | 3384 | 35.52 | 104.00 | 100.57 | 8.73 |
| R5 | 5578 | 58.55 | 116.00 | 116.07 | 3.48 |

**Supplementary Figure 13: Validation of the DT40 wt cell synchronization by elutriation for ATAC-Seq experiments**
Cell cycle analyses by flow cytometry for different fractions of elutriated DT40 wt cells were made after DNA labelling with propidium iodide (PI). The DNA content distributions of cells in G1-phase before (G1) or in G2/M-phase (G2/M) are shown. G1-phase cells are represented in green and G2/M cells in orange. R3, R4 and R5 regions were settled based on the asynchronous cell profile to define the proportion of cells in each fraction. Proportions of cells analyzed for each condition as well as statistics of the distribution of the detected PI values for each region are given in the table below each graph. Black titles on the Y-axis give the name of the cell population analyzed.

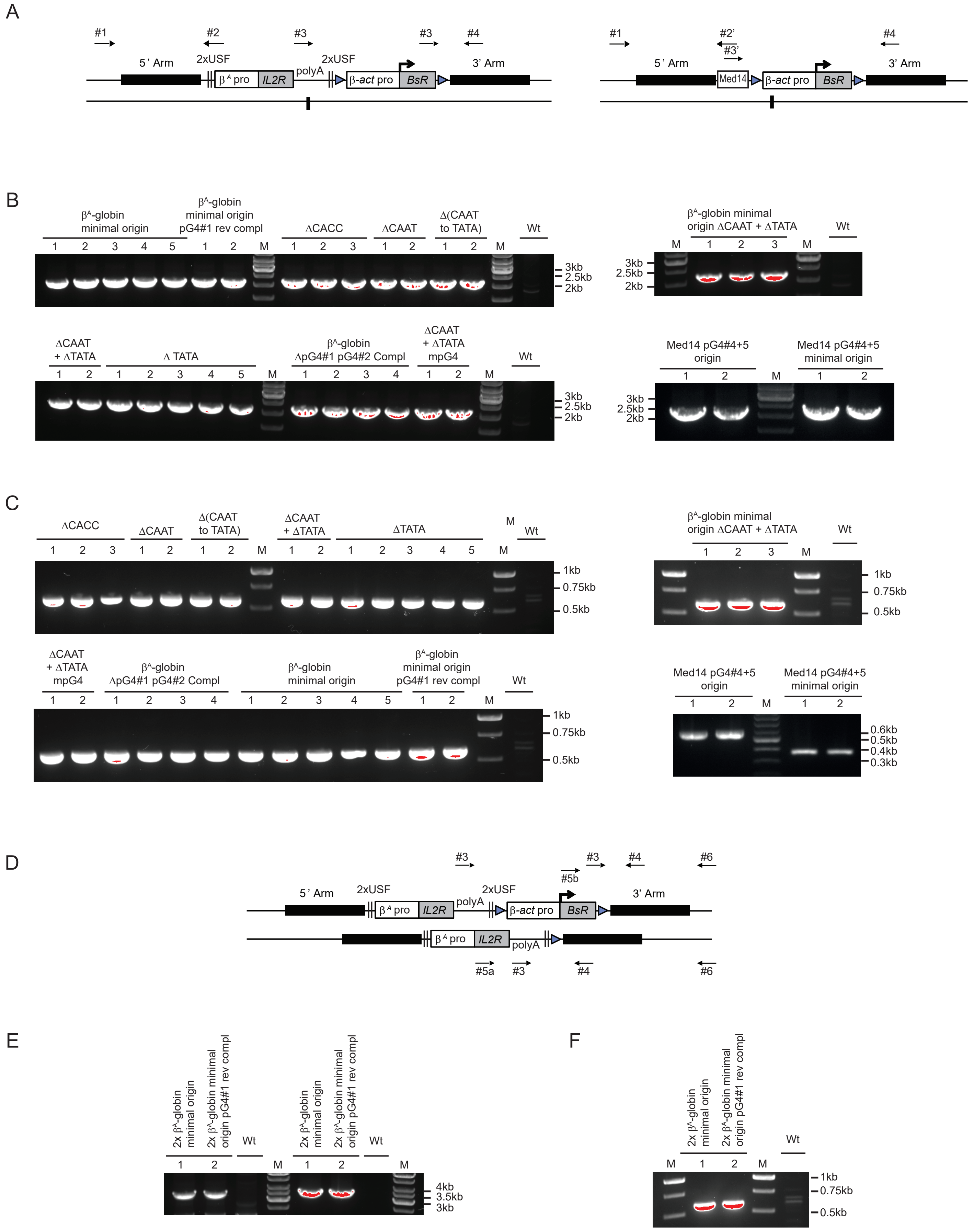


**Supplementary Figure 14: PCR validation of clones selected for homologous recombination**

**Supplementary Figure 14: PCR validation of clones selected for homologous recombination**

(A) Schematic showing the genomic region containing a site-specific integrated construct on one chromosome before excision of the gene of selection (*BsR* gene under the control of the β-actin promoter). Black boxes represent the 5′ and 3' arms of the targeted vector. Positions of primers used for the analysis of correct integration of the constructs by homologous recombination (#1 and #2 or #2') and for the analysis of the correct excision of the *BsR* gene (#3 or #3' and #4) are indicated with arrows. Note that primer #3 can hybridize at two positions since it is located within the polyA signal found 3' of *IL2R* and *BsR* reporter genes. (B, C) PCR products were subjected to electrophoresis in a 1 and 1.5% w/v agarose gel and stained with SYBR safe. The DNA size marker used is a commercial 1 kb plus DNA ladder (M, B) or a 100 bp DNA ladder (M, C). (B) The 2.2 kb PCR products obtained after amplification with primers #1 and #2 and primers #1 and #2' validate the site specific insertion of constructs containing a mutated version of the $\beta^A$ promoter or the Med14 pG4#4+5 origin or the respective minimal origins. The gene of selection was then excised by addition of tamoxifen in clones having made the homologous recombination (see Materials & methods) and the correct excision was controlled by the presence of a 0.6 kb PCR product with primers #3 and #4 and either a 0.35 kb or a 0.5 kb PCR product with primers #3' and #4 . The absence of a correct excision would lead to either a 0.5 kb PCR product in $\beta^A$ globin constructs or no amplification in Med14 constructs (C). (D) Schematic showing the genomic region containing the minimal origin (active or inactive) on both chromosomes before excision of the gene of selection inside the second insertion. To obtain this clone we started from clone 2 for the $\beta^A$-globin minimal active origin and clone 2 for the $\beta^A$-globin minimal inactive origin described in panel B and C. We tested the double insertion with two different primer pairs, the first one #5a and #6 confirmed the presence of the first inserted construct and the second one #5b and #6 validated the insertion of the second construct, before its excision (E). The amplicon between primers #5a and #6 can be obtained only on the chromosome having excised *BsR* since the very high GC content present within the β-actin promoter blocks PCR amplification. We therefore can control whether the second insertion occurred on the same or on the other chromosome. Clones capable of amplifying both 3.5 kb fragments with primer pairs #5a with #6 and #5b with #6 are the one containing the insertion on both chromosomes. Then the gene of selection is excised and the proper excision is tested with primers #3 and #4 (F).

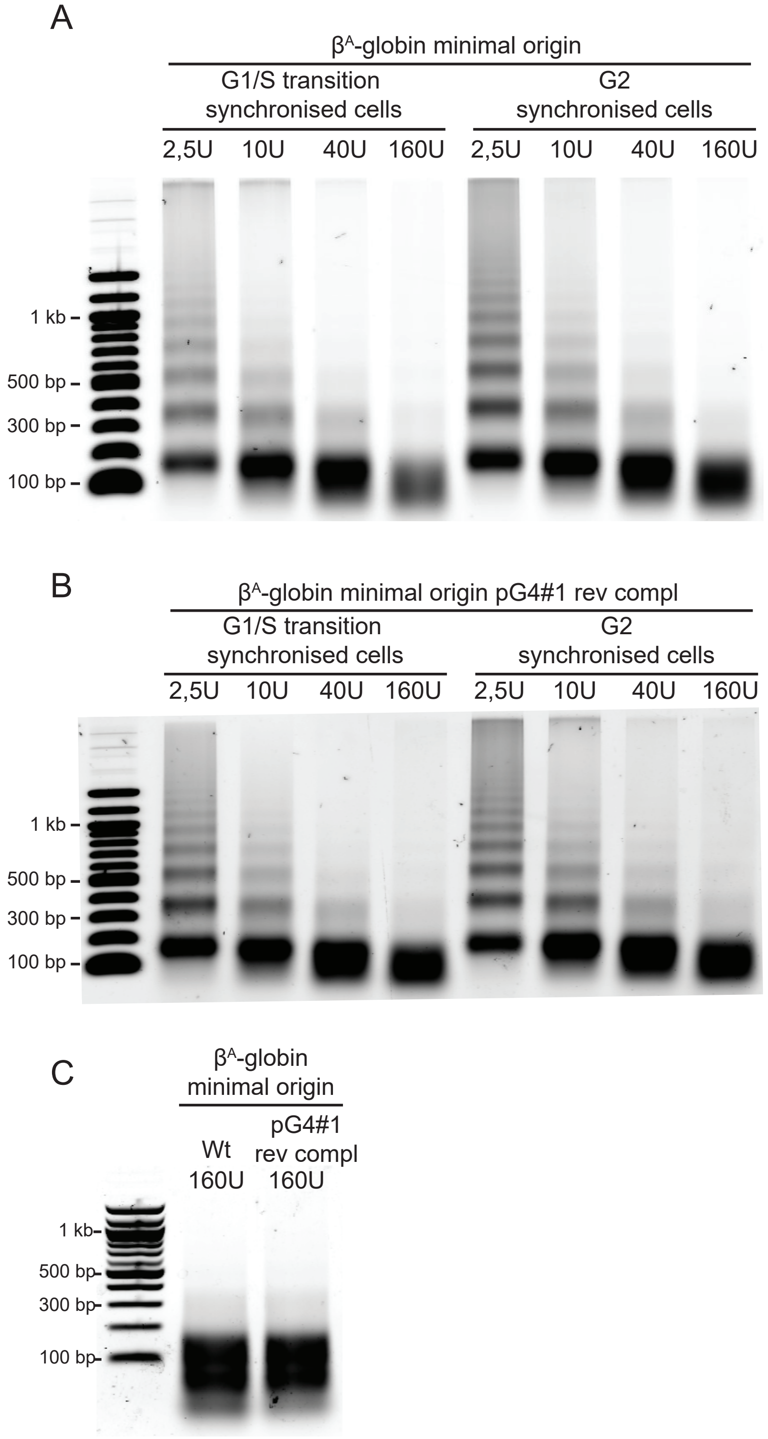


**Supplementary Figure 15: Validation of MNase digestion patterns obtained to analyse nucleosome positioning**
(A-C) Chromatin was extracted from clonal cell lines containing the $\beta^A$-globin minimal origin (A, C) or the $\beta^A$-globin minimal origin containing the pG4#1 reverse complementary sequence (pG4#1 rev compl, B, C). Chromatin was partially digested with exponentially increasing concentrations of micrococcal nuclease (MNAse; 2.5, 10, 40 and 160 U/mL). The four digested DNA samples obtained for each clonal cell line (A and B) or the most digested sample only (160U/mL, C) were subjected to electrophoresis in a 1% w/v agarose gel and stained with SYBR safe. The DNA size marker was a commercial 100bp ladder. (A-B) Digestion patterns of cells synchronised at the G1/S transition are shown on the left and those of cells in G2-phase are shown on the right. (C) Digestion patterns of asynchronous cells are shown.
