## Supplementary Tables for "Dimeric G-quadruplex motifs-induced NFRs determine strong replication origins in vertebrates"

### 1 Logistic regression to quantify genome-wide associations

Here we provide some details regarding the statistical analysis of associations between genomic features. Considering an example with the association of origins with monomeric pG4s (from Supp Table), as illustrated in the contingency table / odds ratio

| All Oris | #inOri | #Ori | %inOri | #inRand | #Rand | %inRand | LogOR |
|---|---|---|---|---|---|---|---|
| Monomeric pG4 | 140,527 | 310,790 | 45 | 90,965 | 306,620 | 30 | 0.7 |

This table contains the probabilities of association that provide the chance of an event happening. If we consider the variable Ori = 1 when a given segment is a replication origins, and Ori = 0 is the segment is a random segment, we can also compute the odds such that:

$$\begin{aligned}\text{Odd of an origin carrying a Mono pG4} &= \frac{140,527}{310,790 - 140,527} = 0.8253525 \\ \text{Odd of a random segment carrying a Mono pG4} &= \frac{90,965}{306,620 - 90,965} = 0.421808\end{aligned}$$

Odds are the ratio of an association vs. non association

$$\text{Odd} = \frac{p}{1-p} = \frac{\text{probability of success}}{\text{probability of failure}}$$

The interest of log-Odds is that they can be used to compare enrichments, for instance here, the enrichment in CGI for origins or for random segments:

$$\begin{aligned}\log \text{Odd}(\text{Ori} = 1, \text{Mono pG4} = 1) &= -0.1919447 \\ \log \text{Odd}(\text{Ori} = 0, \text{Mono pG4} = 1) &= -0.8632051\end{aligned}$$

Here, log-Odds ratio becomes:

$$\log \text{Odd}(\text{Ori} = 1, \text{Mono pG4} = 1) - \log \text{Odd}(\text{Ori} = 0, \text{Mono pG4} = 1) = 0.6712604 > 0$$

Since the logOR is positive, it means that origins are more enriched in Monomeric pG4s than random segments.

This general framework is the logistic regression developed to quantify and test associations between a binary variable and covariates. We used the `emmeans` `R`-package (version 1.7.3) for computing logistic regressions. In this framework, the significance of log-odds ratios is assessed using $z$-tests.

### 2 How to assess if Origins are enriched in Monomeric or in Clustered pG4s ?

In a second step we question whether origins are rather associated with the Monomeric or Clustered Form of pG4s. For this purpose we need to restrict the set of considered origins to origins that actually carry a pG4. Similarly, as a control, we consider random segments that carry a pG4. Then we ask if when origins carry a pG4, they are more enriched in Monomeric pG4 (for instance). Considering the table :

| Oris with pG4 | #inOri | #Ori | %inOri | #inRand | #Rand | %inRand | LogOR |
|---|---|---|---|---|---|---|---|
| Monomeric pG4 | 140,527 | 166,257 | 84 | 90,965 | 103,032 | 88 | -0.3*** |

The computed odds are:

$$\text{Odd of an origin that carries a pG4 of carrying a Mono pG4} = \frac{140,527}{166,257 - 140,527} = 5.461601$$

$$\text{Odd of a random segment that carries a pG4 of carrying a Mono pG4} = \frac{90,965}{103,032 - 90,965} = 7.538328$$

Then the log-Odds ratio becomes:

$$\log \mathrm{Odd}(\mathrm{Ori} = 1, \mathrm{Mono\ pG4} = 1 \mid \mathrm{pG4} = 1) - \log \mathrm{Odd}(\mathrm{Ori} = 0, \mathrm{Mono\ pG4} = 1 \mid \mathrm{pG4} = 1) = -0.3222584,$$

with notation $\mid \mathrm{pG4} = 1$ meaning that the analysis is restricted to segments that carry pG4s. Very interestingly, this logOR has become negative ! meaning that when we restrict the enrichment analysis to segments that carry a pG4, the replication origins are not enriched in the Monomeric form of pG4s, but rather in the Clustered form (logOR=0.5 in Supp. Table). A similar strategy was used to assess the association of H2AZ and NFRs with pG4s.

| pG4 Human | | | | | | | |
|---|---|---|---|---|---|---|---|
| Monomeric pG4 | #inpG4 | #pG4 | %inpG4 | #inRand | #Rand | %inRand | LogOR |
| CGI | 105693 | 1035770 | 10 | 23836 | 1035770 | 2 | 1.6*** |
| TSS | 57046 | 1035770 | 6 | 14442 | 1035770 | 1 | 1.4*** |
| Oris H9 SNS | | | | | | | |
| Ori | 275440 | 1035770 | 27 | 92948 | 1035770 | 9 | 1.3*** |
| Strong Ori | 115157 | 1035770 | 11 | 23479 | 1035770 | 2 | 1.7*** |
| H2AZ | 433256 | 1035770 | 42 | 374299 | 1035770 | 36 | 0.2*** |
| H2AZ in pG4 with Ori | 136311 | 275440 | 50 | 46014 | 92948 | 50 | $0^{ns}$ |
| H2AZ in pG4 without Ori | 296945 | 760330 | 39 | 328285 | 942822 | 35 | 0.2*** |
| chIP G4, Zheng et al. 2020 | 418938 | 1035770 | 40 | 153749 | 1035770 | 15 | 1.4*** |
| chIP G4 in pG4 with Ori | 195993 | 275440 | 71 | 47163 | 92948 | 51 | 0.9*** |
| chIP G4 in pG4 without Ori | 222945 | 760330 | 29 | 106586 | 942822 | 11 | 1.2*** |
| chIP G4, Hansel-Hertsch et al. 2016 | 21599 | 1035770 | 2 | 7020 | 1035770 | 1 | 1.1*** |
| chIP G4 with Ori | 12246 | 275440 | 4 | 2736 | 93023 | 3 | 0.4*** |
| chIP G4 without Ori | 9353 | 760330 | 1 | 4284 | 942747 | 0 | 1*** |
| Core Oris from Akerman et al. (2020), SNS | | | | | | | |
| Ori | 131148 | 1035770 | 13 | 35900 | 1035770 | 4 | 1.4*** |
| Oris EJ30 from Guilbaud et al. (2022), Iniseq2 | | | | | | | |
| Ori | 35706 | 1035770 | 3 | 15335 | 1035770 | 2 | 0.9*** |
| Strong Oris | 7343 | 1035770 | 1 | 4002 | 1035770 | 0 | 0.6*** |
| Clustered pG4 | #inpG4 | #pG4 | %inpG4 | #inRand | #Rand | %inRand | LogOR |
| CGI | 37600 | 156441 | 24 | 3392 | 156441 | 2 | 2.7*** |
| TSS | 21585 | 156441 | 14 | 2130 | 156441 | 1 | 2.5*** |
| Oris H9 SNS | | | | | | | |
| Ori | 73581 | 156441 | 47 | 13646 | 156441 | 9 | 2.2*** |
| Strong Ori | 42713 | 156441 | 27 | 3367 | 156441 | 2 | 2.8*** |
| H2AZ | 67792 | 156441 | 43 | 56299 | 156441 | 36 | 0.3*** |
| H2AZ in pG4 with Ori | 37439 | 73581 | 51 | 6706 | 13646 | 49 | 0.1*** |
| H2AZ in pG4 without Ori | 30353 | 82860 | 37 | 49593 | 142795 | 35 | 0.1*** |
| chIP G4, Zheng et al. 2020 | 106590 | 156441 | 68 | 22782 | 156441 | 15 | 2.5*** |
| chIP G4 in pG4 with Ori | 62188 | 73581 | 84 | 6801 | 13646 | 50 | 1.7*** |
| chIP G4 in pG4 without Ori | 44402 | 82860 | 54 | 15981 | 142795 | 11 | 2.2*** |
| chIP G4, Hansel-Hertsch et al. 2016 | 7042 | 156441 | 4 | 1034 | 156441 | 1 | 2*** |
| chIP G4 with Ori | 4966 | 73581 | 7 | 383 | 13676 | 3 | 0.9*** |
| chIP G4 without Ori | 2076 | 82860 | 2 | 651 | 142765 | 0 | 1.7*** |
| Core Oris from Akerman et al. (2020), SNS | | | | | | | |
| Ori | 50905 | 156441 | 32 | 5265 | 156441 | 3 | 2.6*** |
| Oris EJ30 from Guilbaud et al. (2022), Iniseq2 | | | | | | | |
| Ori | 9542 | 156441 | 6 | 2219 | 156441 | 1 | 1.5*** |
| Strong Oris | 1859 | 156441 | 1 | 579 | 156441 | 0 | 1.2*** |

Table 1: Association of pG4s with genomic features. Associations were computed separately for pG4+ and pG4− and combined. Strong Oris are the top 25% active origins in terms of SNS enrichment. Random pG4 segments were sampled to match the length distribution of observed pG4s (see Material & Methods). #inpG4: number of features in pG4, #pG4: number of pG4s, %inpG4: percentage of features in pG4, #inRand: number of features in random segments, #Rand: number of random segments, %inRand: percentage of feature in random segments, LogOR: log-odd-ratio of the logistic regression to test enrichment in pG4 wrt enrichment in random segments (see Material & Methods).

| pG4 Chicken (DT40) | | | | | | | |
|---|---|---|---|---|---|---|---|
| Monomeric pG4 | #inpG4 | #pG4 | %inpG4 | #inRand | #Rand | %inRand | LogOR |
| CGI | 67003 | 300974 | 22 | 13581 | 300974 | 4 | 1.8*** |
| TSS | 15833 | 300974 | 5 | 3276 | 300974 | 1 | 1.6*** |
| Ori | 92201 | 300974 | 31 | 35127 | 300974 | 12 | 1.2*** |
| Strong Oris | 47142 | 300974 | 16 | 8866 | 300974 | 3 | 1.8*** |
| H2AZ | 196050 | 300974 | 65 | 122209 | 300974 | 41 | 1*** |
| H2AZ in pG4 with Ori | 73530 | 92201 | 80 | 23255 | 35127 | 66 | 0.7*** |
| H2AZ in pG4 without Ori | 122520 | 208773 | 59 | 98954 | 265847 | 37 | 0.9*** |
| NFR in G1 | 28394 | 300974 | 9 | 11446 | 300974 | 4 | 1*** |
| NFR in G1 in pG4 with Ori | 16279 | 92201 | 18 | 3711 | 35127 | 11 | 0.6*** |
| NFR in G1 in pG4 without Ori | 12115 | 208773 | 6 | 7735 | 265847 | 3 | 0.7*** |
| NFR in G2 | 23382 | 300974 | 8 | 8272 | 300974 | 3 | 1.1*** |
| NFR in G2 in pG4 with Ori | 14334 | 92201 | 16 | 3115 | 35127 | 9 | 0.6*** |
| NFR in G2 in pG4 without Ori | 9048 | 208773 | 4 | 5157 | 265847 | 2 | 0.8*** |
| chIP G4, Zheng et al. 2020 | 44677 | 300974 | 15 | 14078 | 300974 | 5 | 1.3*** |
| chIP G4 in pG4 with Ori | 26074 | 92201 | 28 | 5565 | 35127 | 16 | 0.7*** |
| chIP G4 in pG4 without Ori | 18603 | 208773 | 9 | 8513 | 265847 | 3 | 1.1*** |
| Clustered pG4 | #inpG4 | #pG4 | %inpG4 | #inRand | #Rand | %inRand | LogOR |
| CGI | 24593 | 53575 | 46 | 2271 | 53575 | 4 | 3*** |
| TSS | 7158 | 53575 | 13 | 496 | 53575 | 1 | 2.8*** |
| Ori | 27745 | 53575 | 52 | 6279 | 53575 | 12 | 2.1*** |
| Strong Oris | 18425 | 53575 | 34 | 1523 | 53575 | 3 | 2.9*** |
| H2AZ | 42468 | 53575 | 79 | 21595 | 53575 | 40 | 1.7*** |
| H2AZ in pG4 with Ori | 23927 | 27745 | 86 | 4069 | 6279 | 65 | 1.2*** |
| H2AZ in pG4 without Ori | 18541 | 25830 | 72 | 17526 | 47296 | 37 | 1.5*** |
| NFR in G1 | 10181 | 53575 | 19 | 2019 | 53575 | 4 | 1.8*** |
| NFR in G1 in pG4 with Ori | 7186 | 27745 | 26 | 641 | 6279 | 10 | 1.1*** |
| NFR in G1 in pG4 without Ori | 2995 | 25830 | 12 | 1378 | 47296 | 3 | 1.5*** |
| NFR in G2 | 9265 | 53575 | 17 | 1395 | 53575 | 3 | 2.1*** |
| NFR in G2 in pG4 with Ori | 6710 | 27745 | 24 | 523 | 6279 | 8 | 1.3*** |
| NFR in G2 in pG4 without Ori | 2555 | 25830 | 10 | 872 | 47296 | 2 | 1.8*** |
| chIP G4, Zheng et al. 2020 | 16491 | 53575 | 31 | 2450 | 53575 | 5 | 2.2*** |
| chIP G4 in pG4 with Ori | 11444 | 27745 | 41 | 995 | 6279 | 16 | 1.3*** |
| chIP G4 in pG4 without Ori | 5047 | 25830 | 20 | 1455 | 47296 | 3 | 2*** |

| pG4 Mouse (mESC) | | | | | | | |
|---|---|---|---|---|---|---|---|
| Monomeric pG4 | #inpG4 | #pG4 | %inpG4 | #inRand | #Rand | %inRand | LogOR |
| CGI | 51285 | 996347 | 5 | 14481 | 996347 | 2 | 1.3*** |
| TSS | 48146 | 996347 | 5 | 15619 | 996347 | 2 | 1.2*** |
| Ori | 262428 | 996347 | 26 | 137003 | 996347 | 14 | 0.8*** |
| Strong Oris | 105252 | 996347 | 11 | 35297 | 996347 | 4 | 1.2*** |
| H2AZ | 132367 | 996347 | 13 | 49792 | 996347 | 5 | 1.1*** |
| H2AZ in pG4 with Ori | 77739 | 262428 | 30 | 23967 | 137003 | 18 | 0.7*** |
| H2AZ in pG4 without Ori | 54628 | 733919 | 7 | 25825 | 859344 | 3 | 1*** |
| chIP G4, Zheng et al. 2020 | 70078 | 996347 | 7 | 23524 | 996347 | 2 | 1.1*** |
| chIP G4 in pG4 with Ori | 48193 | 262428 | 18 | 13195 | 137003 | 10 | 0.7*** |
| chIP G4 in pG4 without Ori | 21885 | 733919 | 3 | 10329 | 859344 | 1 | 0.9*** |
| Clustered pG4 | #inpG4 | #pG4 | %inpG4 | #inRand | #Rand | %inRand | LogOR |
| CGI | 15610 | 156571 | 10 | 2161 | 156571 | 1 | 2.1*** |
| TSS | 14005 | 156571 | 9 | 2343 | 156571 | 2 | 1.9*** |
| Ori | 54108 | 156571 | 35 | 21070 | 156571 | 14 | 1.2*** |
| Strong Oris | 25881 | 156571 | 16 | 5337 | 156571 | 3 | 1.7*** |
| H2AZ | 26100 | 156571 | 17 | 7711 | 156571 | 5 | 1.4*** |
| H2AZ in pG4 with Ori | 7777 | 102463 | 8 | 4036 | 135501 | 3 | 1*** |
| H2AZ in pG4 without Ori | 18323 | 54108 | 34 | 3675 | 21070 | 17 | 0.9*** |
| chIP G4, Zheng et al. 2020 | 19037 | 156571 | 12 | 3635 | 156571 | 2 | 1.8*** |
| chIP G4 in pG4 with Ori | 14931 | 54108 | 28 | 1977 | 21070 | 9 | 1.3*** |
| chIP G4 in pG4 without Ori | 4106 | 102463 | 4 | 1658 | 135501 | 1 | 1.2*** |

Table 2: Association of pG4s with genomic features. Associations were computed separately for pG4+ and pG4− and combined. Strong Oris are the top 25% active origins in terms of SNS enrichment. Random pG4 segments were sampled to match the length distribution of observed pG4s (see Material & Methods). #inpG4: number of features in pG4, #pG4: number of pG4s, %inpG4: percentage of features in pG4, #inRand: number of features in random segments, #Rand: number of random segments, %inRand: percentage of feature in random segments, LogOR: log-odd-ratio of the logistic regression to test enrichment in pG4 wrt enrichment in random segments (see Material & Methods).

| Oris Human H9 SNS | | | | | | | |
|---|---|---|---|---|---|---|---|
| **All Oris** | **#inOri** | **#Ori** | **%inOri** | **#inRand** | **#Rand** | **%inRand** | **LogOR** |
| CGI | 47,814 | 310,790 | 15 | 16,237 | 290,194 | 6 | 1.1*** |
| TSS | 26,774 | 310,790 | 9 | 7,189 | 290,194 | 2 | 1.3*** |
| all pG4s | 166,257 | 310,790 | 54 | 90,544 | 290,194 | 31 | 0.9*** |
| Monomeric pG4s | 140,527 | 310,790 | 45 | 80,064 | 290,194 | 28 | 0.8*** |
| Clustered pG4s | 53,304 | 310,790 | 17 | 17,958 | 290,194 | 6 | 1.1*** |
| H2AZ | 72,530 | 310,790 | 23 | 39,193 | 290,194 | 14 | 0.6*** |
| chIP G4, Zheng et al. 2020 | 166,857 | 310,790 | 54 | 103,615 | 290,194 | 36 | 0.7*** |
| chIP G4, Hansel-Hertsch et al. 2016 | 9,925 | 310,790 | 3 | 3,466 | 289,256 | 1 | 1*** |
| **Oris with pG4** | **#inOri** | **#Ori** | **%inOri** | **#inRand** | **#Rand** | **%inRand** | **LogOR** |
| CGI | 35,941 | 166,257 | 22 | 9,435 | 90,544 | 10 | 0.9*** |
| TSS | 20,578 | 166,257 | 12 | 4,022 | 90,544 | 4 | 1.1*** |
| Monomeric pG4s | 140,527 | 166,257 | 84 | 80,064 | 90,544 | 88 | -0.3*** |
| Clustered pG4s | 53,304 | 166,257 | 32 | 17,958 | 90,544 | 20 | 0.6*** |
| H2AZ | 42,172 | 166,257 | 25 | 13,176 | 90,544 | 15 | 0.7*** |
| chIP G4, Zheng et al. 2020 | 110,597 | 166,257 | 66 | 49,910 | 90,544 | 55 | 0.5*** |
| chIP G4, Hansel-Hertsch et al. 2016 | 7,136 | 166,257 | 4 | 1,662 | 90,208 | 2 | 0.9*** |

| Oris Human Core from Akerman et al. (2020), SNS | | | | | | | |
|---|---|---|---|---|---|---|---|
| **All Oris** | **#inOri** | **#Ori** | **%inOri** | **#inRand** | **#Rand** | **%inRand** | **LogOR** |
| CGI | 44,494 | 127,248 | 35 | 4,563 | 127,248 | 4 | 2.7*** |
| TSS | 18,843 | 127,248 | 15 | 2,869 | 127,248 | 2 | 2*** |
| all pG4s | 74,716 | 127,248 | 59 | 33,201 | 127,248 | 26 | 1.4*** |
| Monomeric pG4s | 59,854 | 127,248 | 47 | 29,629 | 127,248 | 23 | 1.1*** |
| Clustered pG4s | 33,293 | 127,248 | 26 | 6,350 | 127,248 | 5 | 1.9*** |
| chIP G4, Zheng et al. 2020 | 83,066 | 127,248 | 65 | 36,087 | 127,248 | 28 | 1.6*** |
| chIP G4, Hansel-Hertsch et al. 2016 | 6,036 | 127,248 | 5 | 1,423 | 127,248 | 1 | 1.5*** |
| **Oris with pG4** | **#inOri** | **#Ori** | **%inOri** | **#inRand** | **#Rand** | **%inRand** | **LogOR** |
| CGI | 32,835 | 74,716 | 44 | 2,729 | 33,201 | 8 | 2.2*** |
| TSS | 15,412 | 74,716 | 21 | 1,666 | 33,201 | 5 | 1.6*** |
| Monomeric pG4s | 59,854 | 74,716 | 80 | 29,629 | 33,201 | 89 | -0.7*** |
| Clustered pG4s | 33,293 | 74,716 | 45 | 6,350 | 33,201 | 19 | 1.2*** |
| chIP G4, Zheng et al. 2020 | 58,259 | 74,716 | 78 | 17,723 | 33,201 | 53 | 1.1*** |
| chIP G4, Hansel-Hertsch et al. 2016 | 4,675 | 74,716 | 6 | 726 | 33,339 | 2 | 1.1*** |

| Oris Human EJ30 from Guilbaud et al. (2022), Iniseq2 | | | | | | | |
|---|---|---|---|---|---|---|---|
| **All Oris** | **#inOri** | **#Ori** | **%inOri** | **#inRand** | **#Rand** | **%inRand** | **LogOR** |
| CGI | 30,936 | 47,634 | 65 | 431 | 47,634 | 1 | 5.3*** |
| TSS | 20,390 | 47,634 | 43 | 358 | 47,634 | 1 | 4.6*** |
| all pG4s | 36,827 | 47,634 | 77 | 6,864 | 47,634 | 14 | 3*** |
| Monomeric pG4s | 29,693 | 47,634 | 62 | 6,344 | 47,634 | 13 | 2.4*** |
| Clustered pG4s | 16,396 | 47,634 | 34 | 716 | 47,634 | 2 | 3.5*** |
| chIP G4, Zheng et al. 2020 | 45,763 | 47,634 | 96 | 5,776 | 47,634 | 12 | 5.2*** |
| chIP G4, Hansel-Hertsch et al. 2016 | 7,967 | 47,634 | 17 | 216 | 47,634 | 0 | 3.8*** |
| **Oris with pG4** | **#inOri** | **#Ori** | **%inOri** | **#inRand** | **#Rand** | **%inRand** | **LogOR** |
| CGI | 24,316 | 36,827 | 66 | 174 | 6,864 | 2 | 4.3*** |
| TSS | 15,736 | 36,827 | 43 | 108 | 6,864 | 2 | 3.8*** |
| Monomeric pG4s | 29,693 | 36,827 | 81 | 6,344 | 6,864 | 92 | -1.1*** |
| Clustered pG4s | 16,396 | 36,827 | 44 | 716 | 6,864 | 10 | 1.9*** |
| chIP G4, Zheng et al. 2020 | 35,718 | 36,827 | 97 | 1,655 | 6,864 | 24 | 4.6*** |
| chIP G4, Hansel-Hertsch et al. 2016 | 5,735 | 36,827 | 16 | 66 | 6,983 | 1 | 3*** |

Table 3: Enrichment of Origins in genomic features in human cells. Associations were computed separately 1kb downstream (+ strand) and 1kb upstream the ori peak (− strand), then combined. Oris with pG4s consist in origins that carry a pG4. #inOri: number of features in Oris, #Ori: number of Oris, %inOri: percent of features in Oris, #inRand: number of features in random segments, #Rand: number of random segments, %inRand: percent of features in random segments, LogOR: log-odd-ratio of the logistic regression to test enrichment in Oris with respect to enrichment in random segments (see Material & Methods). When considering origins with pG4s, we restrict the analysis to segments that carry a pG4 (Oris and Random). By doing so we test whether, when origins or random segments are associated with a pG4, the enrichment concerns the monomeric or the Clustered form.

| Oris Chicken DT40, SNS | | | | | | | |
|---|---|---|---|---|---|---|---|
| All Oris | #inOri | #Ori | %inOri | #inRand | #Rand | %inRand | LogOR |
| CGI | 33,233 | 136,136 | 24 | 16,104 | 126,356 | 13 | 0.8*** |
| TSS | 8,681 | 136,136 | 6 | 2,180 | 126,356 | 2 | 1.4*** |
| all pG4s | 57,209 | 136,136 | 42 | 40,495 | 126,356 | 32 | 0.4*** |
| Monomeric pG4s | 48,259 | 136,136 | 35 | 35,981 | 126,356 | 28 | 0.3*** |
| Clustered pG4s | 19,412 | 136,136 | 14 | 8,868 | 126,610 | 7 | 0.8*** |
| H2AZ | 84,841 | 136,136 | 62 | 67,347 | 126,356 | 53 | 0.4*** |
| NFR in G1 | 16,035 | 136,136 | 12 | 6,626 | 126,356 | 5 | 0.9*** |
| NFR in G2 | 13,560 | 136,136 | 10 | 4,421 | 126,356 | 4 | 1.1*** |
| chIP G4, Zheng et al. 2020 | 24,176 | 136,136 | 18 | 9,321 | 126,356 | 7 | 1*** |
| Oris with pG4 | #inOri | #Ori | %inOri | #inRand | #Rand | %inRand | LogOR |
| CGI | 22,736 | 57,209 | 40 | 10,125 | 40,495 | 25 | 0.7*** |
| TSS | 6,480 | 57,209 | 11 | 1,508 | 40,495 | 4 | 1.2*** |
| Monomeric pG4s | 48,259 | 57,209 | 84 | 35,981 | 40,495 | 89 | -0.4*** |
| Clustered pG4s | 19,412 | 57,209 | 34 | 8,694 | 40,495 | 22 | 0.6*** |
| H2AZ | 41,959 | 57,209 | 73 | 26,393 | 40,495 | 65 | 0.4*** |
| NFR in G1 | 10,049 | 57,209 | 18 | 3,240 | 40,495 | 8 | 0.9*** |
| NFR in G2 | 8,960 | 57,209 | 16 | 2,234 | 40,495 | 6 | 1.2*** |
| chIP G4, Zheng et al. 2020 | 15,669 | 57,209 | 27 | 4,937 | 40,495 | 12 | 1*** |

| Oris Mouse mESC, SNS | | | | | | | |
|---|---|---|---|---|---|---|---|
| All Oris | #inOri | #Ori | %inOri | #inRand | #Rand | %inRand | LogOR |
| CGI | 36,889 | 411,762 | 9 | 2,294 | 190,458 | 1 | 2.1*** |
| TSS | 32,713 | 411,762 | 8 | 3,376 | 190,458 | 2 | 1.6*** |
| all pG4s | 176,495 | 411,762 | 43 | 56,545 | 190,458 | 30 | 0.6*** |
| Monomeric pG4s | 147,902 | 411,762 | 36 | 50,524 | 190,458 | 26 | 0.4*** |
| Clustered pG4s | 44,206 | 411,762 | 11 | 8,257 | 190,458 | 4 | 1*** |
| H2AZ | 80,454 | 411,762 | 20 | 21,795 | 190,458 | 11 | 0.6*** |
| chIP G4, Zheng et al. 2020 | 45,480 | 411,762 | 11 | 7,015 | 190,458 | 4 | 1.2*** |
| Oris with pG4 | #inOri | #Ori | %inOri | #inRand | #Rand | %inRand | LogOR |
| CGI | 24,309 | 176,495 | 14 | 1,123 | 56,545 | 2 | 2.1*** |
| TSS | 21,383 | 176,495 | 12 | 1,541 | 56,545 | 3 | 1.6*** |
| Monomeric pG4s | 147,902 | 176,495 | 84 | 50,524 | 56,545 | 89 | -0.5*** |
| Clustered pG4s | 44,206 | 176,495 | 25 | 8,257 | 56,545 | 15 | 0.7*** |
| H2AZ | 46,300 | 176,495 | 26 | 8,894 | 56,545 | 16 | 0.6*** |
| chIP G4, Zheng et al. 2020 | 29,084 | 176,495 | 16 | 3,054 | 56,545 | 5 | 1.2*** |

Table 4: Enrichment of Origins in genomic features in chicken and mouse cells. Associations were computed separately 1kb downstream (+ strand) and 1kb upstream the ori peak (− strand), then combined. Oris with pG4s consist in origins that carry a pG4. #inOri: number of features in Oris, #Ori: number of Oris, %inOri: percent of features in Oris, #inRand: number of features in random segments, #Rand: number of random segments, %inRand: percent of features in random segments, LogOR: log-odd-ratio of the logistic regression to test enrichment in Oris with respect to enrichment in random segments (see Material & Methods). When considering origins with pG4s, we restrict the analysis to segments that carry a pG4 (Oris and Random). By doing so we test whether, when origins or random segments are associated with a pG4, the enrichment concerns the monomeric or the Clustered form.

| Strong Oris Human H9 SNS | | | | | | | |
|---|---|---|---|---|---|---|---|
| All Oris | #inOri | #Ori | %inOri | #inRand | #Rand | %inRand | LogOR |
| CGI | 24,786 | 77,698 | 32 | 4,462 | 72,548 | 6 | 2*** |
| TSS | 12,147 | 77,698 | 16 | 2,056 | 72,548 | 3 | 1.8*** |
| all pG4s | 60,260 | 77,698 | 78 | 23,554 | 72,548 | 32 | 2*** |
| Monomeric pG4s | 48,374 | 77,698 | 62 | 20,940 | 72,548 | 29 | 1.4*** |
| Clustered pG4s | 29,234 | 77,698 | 38 | 4,499 | 72,548 | 6 | 2.2*** |
| H2AZ | 20,614 | 77,698 | 26 | 9,924 | 72,548 | 14 | 0.7*** |
| chIP G4, Zheng et al. 2020 | 67,232 | 77,698 | 86 | 26,380 | 72,548 | 36 | 2.4*** |
| chIP G4, Hansel-Hertsch et al. 2016 | 4,009 | 77,698 | 5 | 997 | 72,314 | 1 | 1.4*** |
| Oris with pG4 | #inOri | #Ori | %inOri | #inRand | #Rand | %inRand | LogOR |
| CGI | 20,600 | 60,260 | 34 | 2,625 | 23,554 | 11 | 1.4*** |
| TSS | 10,480 | 60,260 | 17 | 1,179 | 23,554 | 5 | 1.4*** |
| Monomeric pG4s | 48,374 | 60,260 | 80 | 20,940 | 23,554 | 89 | -0.7*** |
| Clustered pG4s | 29,234 | 60,260 | 48 | 4,499 | 23,554 | 19 | 1.4*** |
| H2AZ | 16,569 | 60,260 | 28 | 3,520 | 23,554 | 15 | 0.8*** |
| chIP G4, Zheng et al. 2020 | 53,823 | 60,260 | 89 | 12,974 | 23,554 | 55 | 1.9*** |
| chIP G4, Hansel-Hertsch et al. 2016 | 3,339 | 60,260 | 6 | 493 | 23,177 | 2 | 1*** |

| Strong Oris Human EJ30 from Guilbaud et al. (2022), Iniseq2 | | | | | | | |
|---|---|---|---|---|---|---|---|
| All Oris | #inOri | #Ori | %inOri | #inRand | #Rand | %inRand | LogOR |
| CGI | 11,494 | 11,906 | 96 | 121 | 11,908 | 1 | 7.9*** |
| TSS | 8,886 | 11,906 | 75 | 85 | 11,908 | 1 | 6*** |
| all pG4s | 9,673 | 11,906 | 81 | 1,821 | 11,908 | 15 | 3.2*** |
| Monomeric pG4s | 7,644 | 11,906 | 64 | 1,682 | 11,908 | 14 | 2.4*** |
| Clustered pG4s | 4,626 | 11,906 | 39 | 190 | 11,908 | 2 | 3.7*** |
| chIP G4, Zheng et al. 2020 | 11,719 | 11,906 | 98 | 1,530 | 11,908 | 13 | 6.1*** |
| chIP G4, Hansel-Hertsch et al. 2016 | 3,218 | 11,906 | 27 | 68 | 11,908 | 1 | 4.2*** |
| Oris with pG4 | #inOri | #Ori | %inOri | #inRand | #Rand | %inRand | LogOR |
| CGI | 9,384 | 9,673 | 97 | 37 | 1,821 | 2 | 7.4*** |
| TSS | 7,234 | 9,673 | 75 | 20 | 1,821 | 1 | 5.6*** |
| Monomeric pG4s | 7,644 | 9,673 | 79 | 1,682 | 1,821 | 92 | -1.2*** |
| Clustered pG4s | 4,626 | 9,673 | 48 | 190 | 1,821 | 10 | 2.1*** |
| chIP G4, Zheng et al. 2020 | 9,543 | 9,673 | 99 | 462 | 1,821 | 25 | 5.4*** |
| chIP G4, Hansel-Hertsch et al. 2016 | 2,551 | 9,673 | 26 | 16 | 1,838 | 1 | 3.7*** |

Table 5: Enrichment of Strong Origins in genomic features in human cells. Strong origins: top 25% most active origins in terms of SNS enrichment. Associations were computed separately 1kb downstream (+ strand) and 1kb upstream the ori peak (− strand), then combined. Strong Oris with pG4s consist in strong origins that carry a pG4. #inOri: number of features in Strong Oris, #Ori: number of Strong Oris, %inOri: percent of features in Strong Oris, #inRand: number of features in random segments, #Rand: number of random segments, %inRand: percent of features in random segments, LogOR: log-odd-ratio of the logistic regression to test enrichment in Strong Oris with respect to enrichment in random segments (see Material & Methods). When considering strong origins with pG4s, we restrict the analysis to segments that carry a pG4 (Strong Oris and Random). By doing so we test whether, when strong origins or random segments are associated with a pG4, the enrichment concerns the monomeric or the Clustered form.

| Strong Oris Chicken DT40, SNS | | | | | | | |
|---|---|---|---|---|---|---|---|
| All Oris | #inOri | #Ori | %inOri | #inRand | #Rand | %inRand | LogOR |
| CGI | 16,085 | 34,034 | 47 | 3,995 | 31,589 | 13 | 1.8*** |
| TSS | 3,482 | 34,034 | 10 | 566 | 31,589 | 2 | 1.8*** |
| all pG4s | 25,746 | 34,034 | 76 | 9,895 | 31,589 | 31 | 1.9*** |
| Monomeric pG4s | 20,909 | 34,034 | 61 | 8,866 | 31,589 | 28 | 1.4*** |
| Clustered pG4s | 12,232 | 34,034 | 36 | 2,064 | 31,652 | 6 | 2.1*** |
| H2AZ | 24,968 | 34,034 | 73 | 17,302 | 31,589 | 55 | 0.8*** |
| NFR in G1 | 5,864 | 34,034 | 17 | 1,755 | 31,589 | 6 | 1.3*** |
| NFR in G2 | 4,953 | 34,034 | 15 | 1,157 | 31,589 | 4 | 1.5*** |
| chIP G4, Zheng et al. 2020 | 9,576 | 34,034 | 28 | 2,487 | 31,589 | 8 | 1.5*** |
| Oris with pG4 | #inOri | #Ori | %inOri | #inRand | #Rand | %inRand | LogOR |
| CGI | 13,127 | 25,746 | 51 | 2,511 | 9,895 | 25 | 1.1*** |
| TSS | 2,959 | 25,746 | 12 | 403 | 9,895 | 4 | 1.1*** |
| Monomeric pG4s | 20,909 | 25,746 | 81 | 8,866 | 9,895 | 90 | -0.7*** |
| Clustered pG4s | 12,232 | 25,746 | 48 | 1,999 | 9,895 | 20 | 1.3*** |
| H2AZ | 19,381 | 25,746 | 75 | 6,488 | 9,895 | 66 | 0.5*** |
| NFR in G1 | 4,668 | 25,746 | 18 | 861 | 9,895 | 9 | 0.8*** |
| NFR in G2 | 4,063 | 25,746 | 16 | 589 | 9,895 | 6 | 1.1*** |
| chIP G4, Zheng et al. 2020 | 7,664 | 25,746 | 30 | 1,248 | 9,895 | 13 | 1.1*** |

| Strong Oris Mouse mESC, SNS | | | | | | | |
|---|---|---|---|---|---|---|---|
| All Oris | #inOri | #Ori | %inOri | #inRand | #Rand | %inRand | LogOR |
| CGI | 26,816 | 102,696 | 26 | 627 | 47,614 | 1 | 3.3*** |
| TSS | 21,519 | 102,696 | 21 | 888 | 47,614 | 2 | 2.6*** |
| all pG4s | 60,685 | 102,696 | 59 | 13,903 | 47,614 | 29 | 1.3*** |
| Monomeric pG4s | 50,942 | 102,696 | 50 | 12,438 | 47,614 | 26 | 1*** |
| Clustered pG4s | 18,605 | 102,696 | 18 | 1,986 | 47,614 | 4 | 1.6*** |
| H2AZ | 38,409 | 102,696 | 37 | 5,140 | 47,614 | 11 | 1.6*** |
| chIP G4, Zheng et al. 2020 | 28,935 | 102,696 | 28 | 1,901 | 47,614 | 4 | 2.2*** |
| Oris with pG4 | #inOri | #Ori | %inOri | #inRand | #Rand | %inRand | LogOR |
| CGI | 18,529 | 60,685 | 30 | 299 | 13,903 | 2 | 3*** |
| TSS | 15,300 | 60,685 | 25 | 383 | 13,903 | 3 | 2.5*** |
| Monomeric pG4s | 50,942 | 60,685 | 84 | 12,438 | 13,903 | 90 | -0.5*** |
| Clustered pG4s | 18,605 | 60,685 | 31 | 1,986 | 13,903 | 14 | 1*** |
| H2AZ | 25,730 | 60,685 | 42 | 2,006 | 13,903 | 14 | 1.5*** |
| chIP G4, Zheng et al. 2020 | 20,124 | 60,685 | 33 | 830 | 13,903 | 6 | 2.1*** |

Table 6: Enrichment of Strong Origins in genomic features in chicken and mouse cells. Strong origins: top 25% most active origins in terms of SNS enrichment. Associations were computed separately 1kb downstream (+ strand) and 1kb upstream the ori peak (− strand), then combined. Strong Oris with pG4s consist in strong origins that carry a pG4. #inOri: number of features in Strong Oris, #Ori: number of Strong Oris, %inOri: percent of features in Strong Oris, #inRand: number of features in random segments, #Rand: number of random segments, %inRand: percent of features in random segments, LogOR: log-odd-ratio of the logistic regression to test enrichment in Strong Oris with respect to enrichment in random segments (see Material & Methods). When considering strong origins with pG4s, we restrict the analysis to segments that carry a pG4 (Strong Oris and Random). By doing so we test whether, when strong origins or random segments are associated with a pG4, the enrichment concerns the monomeric or the Clustered form.

| H2AZ Human H9 | | | | | | | |
|---|---|---|---|---|---|---|---|
| All H2AZ | #inH2AZ | #H2AZ | %inH2AZ | #inRand | #Rand | %inRand | LogOR |
| CGI | 48,517 | 440,058 | 11 | 8,140 | 706,920 | 1 | 2.4*** |
| TSS | 35,253 | 440,058 | 8 | 4,309 | 706,920 | 1 | 2.7*** |
| Oris | 81,497 | 440,058 | 18 | 52,589 | 706,920 | 7 | 1*** |
| Strong Oris | 26,295 | 440,058 | 6 | 12,304 | 706,920 | 2 | 1.3*** |
| Monomeric pG4s | 92,474 | 440,058 | 21 | 94,468 | 706,920 | 13 | 0.5*** |
| Clustered pG4 | 22,924 | 440,058 | 5 | 15,414 | 706,920 | 2 | 0.9*** |
| chIP G4, Zheng et al. 2020 | 120,845 | 440,058 | 28 | 92,154 | 706,920 | 13 | 0.9*** |
| chIP G4, Hansel-Hertsch et al. 2016 | 17,006 | 440,058 | 4 | 2,074 | 706,920 | 0 | 2.6*** |
| H2AZ with pG4 | #inH2AZ | #H2AZ | %inH2AZ | #inRand | #Rand | %inRand | LogOR |
| CGI | 31,486 | 104,376 | 30 | 5,081 | 104,032 | 5 | 2.1*** |
| TSS | 22,261 | 104,376 | 21 | 1,827 | 104,032 | 2 | 2.7*** |
| Oris | 43,335 | 104,376 | 42 | 23,101 | 104,032 | 22 | 0.9*** |
| Strong Oris | 19,378 | 104,376 | 19 | 8,383 | 104,032 | 8 | 1*** |
| Monomeric pG4s | 92,474 | 104,376 | 89 | 94,468 | 104,032 | 91 | -0.2*** |
| Clustered pG4 | 22,924 | 104,376 | 22 | 15,414 | 104,032 | 15 | 0.5*** |
| chIP G4, Zheng et al. 2020 | 59,400 | 104,376 | 57 | 35,933 | 104,032 | 34 | 0.9*** |
| chIP G4, Hansel-Hertsch et al. 2016 | 9,458 | 104,376 | 9 | 741 | 104,032 | 1 | 2.6*** |

| H2AZ Chicken DT40 | | | | | | | |
|---|---|---|---|---|---|---|---|
| All H2AZ | #inH2AZ | #H2AZ | %inH2AZ | #inRand | #Rand | %inRand | LogOR |
| CGI | 55,537 | 539,658 | 10 | 9,087 | 588,604 | 2 | 2*** |
| TSS | 14,730 | 539,658 | 3 | 1,313 | 588,604 | 0 | 2.5*** |
| Oris | 114,184 | 539,658 | 21 | 42,588 | 588,604 | 7 | 1.2*** |
| Strong Oris | 35,820 | 539,658 | 7 | 7,557 | 588,604 | 1 | 1.7*** |
| Monomeric pG4s | 117,530 | 539,658 | 22 | 48,331 | 588,604 | 8 | 1.1*** |
| Clustered pG4 | 30,601 | 539,658 | 6 | 6,089 | 588,604 | 1 | 1.7*** |
| NFR in G1 | 41,200 | 539,658 | 8 | 11,067 | 588,604 | 2 | 1.5*** |
| chIP G4, Zheng et al. 2020 | 51,859 | 539,658 | 10 | 13,286 | 588,604 | 2 | 1.5*** |
| H2AZ with pG4 | #inH2AZ | #H2AZ | %inH2AZ | #inRand | #Rand | %inRand | LogOR |
| CGI | 35,204 | 133,606 | 26 | 4,753 | 52,195 | 9 | 1.3*** |
| TSS | 9,622 | 133,606 | 7 | 526 | 52,195 | 1 | 2*** |
| Oris | 50,028 | 133,606 | 37 | 10,148 | 52,195 | 19 | 0.9*** |
| Strong Oris | 24,724 | 133,606 | 18 | 4,329 | 52,195 | 8 | 0.9*** |
| Monomeric pG4s | 117,530 | 133,606 | 88 | 48,331 | 52,195 | 93 | -0.5*** |
| Clustered pG4 | 30,601 | 133,606 | 23 | 6,089 | 52,195 | 12 | 0.8*** |
| NFR in G1 | 16,356 | 133,606 | 12 | 1,545 | 52,195 | 3 | 1.5*** |
| chIP G4, Zheng et al. 2020 | 25,257 | 133,606 | 19 | 2,872 | 52,195 | 6 | 1.4*** |

| H2AZ Mouse mESC | | | | | | | |
|---|---|---|---|---|---|---|---|
| All H2AZ | #inH2AZ | #H2AZ | %inH2AZ | #inRand | #Rand | %inRand | LogOR |
| CGI | 31,504 | 156,718 | 20 | 2,006 | 150,554 | 1 | 2.9*** |
| TSS | 28,834 | 156,718 | 18 | 2,588 | 150,554 | 2 | 2.6*** |
| Oris | 80,498 | 156,718 | 51 | 34,420 | 150,554 | 23 | 1.3*** |
| Strong Oris | 42,588 | 156,718 | 27 | 10,959 | 150,554 | 7 | 1.6*** |
| Monomeric pG4s | 61,985 | 156,718 | 40 | 38,099 | 150,554 | 25 | 0.7*** |
| Clustered pG4 | 16,025 | 156,718 | 10 | 7,082 | 150,554 | 5 | 0.8*** |
| chIP G4, Zheng et al. 2020 | 36,139 | 156,718 | 23 | 5,304 | 150,554 | 4 | 2.1*** |
| H2AZ with pG4 | #inH2AZ | #H2AZ | %inH2AZ | #inRand | #Rand | %inRand | LogOR |
| CGI | 19,924 | 70,898 | 28 | 1,026 | 43,146 | 2 | 2.8*** |
| TSS | 17,618 | 70,898 | 25 | 1,215 | 43,146 | 3 | 2.4*** |
| Oris | 43,058 | 70,898 | 61 | 13,663 | 43,146 | 32 | 1.2*** |
| Strong Oris | 26,635 | 70,898 | 38 | 5,448 | 43,146 | 13 | 1.4*** |
| Monomeric pG4s | 61,985 | 70,898 | 87 | 38,099 | 43,146 | 88 | -0.1*** |
| Clustered pG4 | 16,025 | 70,898 | 23 | 7,082 | 43,146 | 16 | 0.4*** |
| chIP G4, Zheng et al. 2020 | 21,675 | 70,898 | 31 | 2,549 | 43,146 | 6 | 1.9*** |

Table 7: Enrichment of H2AZ peaks in genomic features. Associations were computed separately 1kb downstream (+ strand) and 1kb upstream the H2AZ peak (− strand), then combined. H2AZ with pG4s consist in H2AZ that carry a pG4. #inH2AZ: number of features in H2AZ, #H2AZ: number of H2AZ, %inH2AZ: percent of features in H2AZ, #inRand: number of features in random segments, #Rand: number of random segments, %inRand: percent of features in random segments, LogOR: log-odd-ratio of the logistic regression to test enrichment in H2AZ with respect to enrichment in random segments (see Material & Methods). When considering H2AZ with pG4s, we restrict the analysis to segments that carry a pG4 (H2AZ and Random). By doing so we test whether, when H2AZ or random segments are associated with a pG4, the enrichment concerns the monomeric or the clustered form.

| NFR Chicken DT40 | | | | | | | |
|---|---|---|---|---|---|---|---|
| All NFR | #inNFR | #NFR | %inNFR | #inRand | #Rand | %inRand | LogOR |
| CGI | 15,808 | 46,256 | 34 | 7,163 | 46,256 | 16 | 1*** |
| TSS | 7,581 | 46,256 | 16 | 943 | 46,256 | 2 | 2.2*** |
| Oris | 17,015 | 46,256 | 37 | 10,492 | 46,256 | 23 | 0.7*** |
| Strong Oris | 7,274 | 46,256 | 16 | 5,550 | 46,256 | 12 | 0.3*** |
| All pG4s | 16,803 | 46,256 | 36 | 15,158 | 46,256 | 33 | 0.2*** |
| Monomeric pG4s | 13,918 | 46,256 | 30 | 13,136 | 46,256 | 28 | 0.1*** |
| Clustered pG4s | 6,157 | 46,256 | 13 | 4,310 | 46,256 | 9 | 0.4*** |
| H2AZ | 33,954 | 46,256 | 73 | 23,327 | 46,256 | 50 | 1*** |
| NFR with pG4 | #inNFR | #NFR | %inNFR | #inRand | #Rand | %inRand | LogOR |
| CGI | 10,259 | 16,803 | 61 | 4,920 | 15,158 | 32 | 1.2*** |
| TSS | 5,119 | 16,803 | 30 | 658 | 15,158 | 4 | 2.3*** |
| Oris | 9,932 | 16,803 | 59 | 5,976 | 15,158 | 39 | 0.8*** |
| Strong Oris | 5,218 | 16,803 | 31 | 4,217 | 15,158 | 28 | 0.2*** |
| Clustered pG4s | 6,157 | 16,803 | 37 | 4,310 | 15,158 | 28 | 0.4*** |
| Monomeric pG4s | 13,918 | 16,803 | 83 | 13,136 | 15,158 | 87 | -0.3*** |
| H2AZ | 14,961 | 16,803 | 89 | 10,045 | 15,158 | 66 | 1.4*** |

Table 8: Enrichment of NFR (in G1) in genomic features. Associations were computed separately 1kb downstream (+ strand) and 1kb upstream the NFR (− strand), then combined. NFR with pG4s consist in NFRs that carry a pG4. #inNFR: number of features in NFR, #NFR: number of NFRs, %inNFR: percent of features in NFR, #inRand: number of features in random segments, #Rand: number of random segments, %inRand: percent of features in random segments, LogOR: log-odd-ratio of the logistic regression to test enrichment in NFRs with respect to enrichment in random segments (see Material & Methods). When considering NFRs with pG4s, we restrict the analysis to segments that carry a pG4 (NFRs and Random). By doing so we test whether, when NFRs or random segments are associated with a pG4, the enrichment concerns the monomeric or the clustered form.

| | Forward primer sequence | Reverse primer sequence | Genomic position (Build Mars 2018) |
|---|---|---|---|
| PBNs enrichment analysis | | | |
| Primers 0 | GGGCTATTCAGCTTGTCTAG | GCCACCTCAACTTTTGTATAC | |
| Primers 1 | GGGGACTGCTCACGTTCATCA | AATGTGGCGTGTGGGATCTC | |
| Primers 1' | CTACACAGAGGTCCTGCTG | GTGAAGAGAAGCCTCAGGCA | |
| Primers 2 | GGGAGCAAGAGCCCAGAC | GTGAGCAGTCCCCACATCAG | |
| Primers 3 | GGGCTATTCAGCTTGTCTAG | TGGAACGTAACTGCGCAACT | |
| Primers 4 | GGGCTATTCAGCTTGTCTAG | GTATACAAAGTTGCTCTTGTG | |
| Bkgd | TCCATACAGCCACAACAGCA | TGTGGAAGAGTTTCAGTCCAGG | chr1:72804257+72804372 |
| ρ-globin | GACGGTCAGGTTTGCCAAAG | TCCTGAGGATACGTTTTTCAG | chr1:197287850-197288114 |
| Replication timing analysis | | | |
| Early | GACGGTCAGGTTTGCCAAAG | TCCTGAGGATACGTTTTTCAG | chr1:197287850-197288114 |
| With | GGGGACTGCTCACGTTCATCA | AATGTGGCGTGTGGGATCTC | |
| Both | TCCATACAGCCACAACAGCA | TGTGGAAGAGTTTCAGTCCAGG | chr1:72804257+72804372 |
| Without | CAGGACAGCAGGTATTCACA | GGCCTGAACACTGTGTCAAT | chr1:72798802+72798956 |
| Screening of targeted integration | | | |
| 5'-screening-site (primers 1 and 2) | GTGCAGCATCAGTGGATAAAGT | GCCACCTCAACTTTTGTATAC | |
| 5'-screening-site (primers 1 and 2') | GTGCAGCATCAGTGGATAAAGT | TGGAACGTAACTGCGCAACT | |
| 3'-screening-site (primers 5a and 6) allele 1 | CTACACAGAGGTCCTGCTG | CCACATGTTTATTGCATACGGC | |
| 3'-screening-site (primers 5b and 6) allele 2 | CGGCAGTACATATTGAAGCGT | CCACATGTTTATTGCATACGGC | |
| Screening of site specific excision | | | |
| BsR cassette excision (primers 3 and 4) | CCCCCTGAACCTGAAACATAA | CCAGGCTGTACTCTGAATCATCT | |
| BsR cassette excision (primers 3' and 4) | TGTATACAAAAGTTGCGCAGTTACGTTCCA | CCAGGCTGTACTCTGAATCATCT | |
| Copy number quantification | | | |
| With ($\beta^A$-globin constructs) | GGGGACTGCTCACGTTCATCA | AATGTGGCGTGTGGGATCTC | |
| With (Med14 constructs) | GGGCTATTCAGCTTGTCTAG | TGGAACGTAACTGCGCAACT | |
| Both | TCCATACAGCCACAACAGCA | TGTGGAAGAGTTTCAGTCCAGG | chr1:72804257+72804372 |
| RNA quantification | | | |
| *Med14* gene | TGGGCTAATAATGCTGGAAAGGT | TAGAGAAGCCAGACGATCAGCA | chr1:113541601+113542342 |
| *Il2R* gene | GGGGACTGCTCACGTTCATCA | AATGTGGCGTGTGGGATCTC | |
| Primers 1 and 2 | GGGCTATTCAGCTTGTCTAG | AATGTGGCGTGTGGGATCTC | |

**Table 9: Primer sets used for quantitative PCR**

| | With concentration | Both concentration | Ratio With/Both |
|---|---|---|---|
| ΔCAAT #1 | 3.49 | 6.79 | 0.51 |
| ΔCAAT #2 | 3.51 | 7.01 | 0.50 |
| ΔTATA #1 | 4.04 | 9.02 | 0.45 |
| ΔTATA #2 | 3.83 | 7.75 | 0.49 |
| ΔTATA #3 | 7.98 | 18.50 | 0.43 |
| ΔTATA #4 | 6.28 | 14.57 | 0.43 |
| ΔTATA #5 | 4.86 | 11.30 | 0.43 |
| ΔCAAT+ΔTATA #1 | 0.49 | 0.97 | 0.50 |
| ΔCAAT+ΔTATA #2 | 3.68 | 7.95 | 0.46 |
| ΔCACC #1 | 3.61 | 7.25 | 0.50 |
| ΔCACC #2 | 3.75 | 7.22 | 0.52 |
| ΔCACC #3 | 6.13 | 13.60 | 0.45 |
| Δ(CAAT to TATA) #1 | 3.33 | 7.02 | 0.48 |
| Δ(CAAT to TATA) #2 | 3.36 | 6.91 | 0.49 |
| ΔCAAT+ΔTATA mpG4 #1 | 0.31 | 0.57 | 0.55 |
| ΔCAAT+ΔTATA mpG4 #2 | 0.44 | 0.75 | 0.58 |
| $\beta^A$ -globin minimal origin #1 | 7.15 | 15.00 | 0.48 |
| $\beta^A$ -globin minimal origin #2 | 0.76 | 1.17 | 0.65 |
| $\beta^A$ -globin minimal origin #3 | 0.40 | 0.75 | 0.54 |
| $\beta^A$ -globin minimal origin #4 | 6.80 | 15.27 | 0.45 |
| $\beta^A$ -globin minimal origin #5 | 6.25 | 13.97 | 0.45 |
| $\beta^A$ -globin minimal origin pG4#1 rev compl #1 | 4.02 | 6.72 | 0.60 |
| $\beta^A$ -globin minimal origin pG4#1 rev compl #2 | 0.49 | 0.97 | 0.50 |
| 2x $\beta^A$-globin minimal origin #1 | 0.51 | 0.63 | 0.81 |
| 2x $\beta^A$ -globin minimal origin pG4#1 rev compl #1 | 0.35 | 0.37 | 0.92 |
| ΔpG4#1 pG4#2 Compl #1 | 0.32 | 0.56 | 0.57 |
| ΔpG4#1 pG4#2 Compl #2 | 6.60 | 14.87 | 0.44 |
| ΔpG4#1 pG4#2 Compl #3 | 0.30 | 0.53 | 0.56 |
| ΔpG4#1 pG4#2 Compl #4 | 5.50 | 12.80 | 0.43 |
| $\beta^A$ -globin minimal origin ΔCAAT+ ΔTATA #1 | 0.57 | 1.09 | 0.52 |
| βA -globin minimal origin ΔCAAT+ ΔTATA #2 | 0.50 | 0.91 | 0.55 |
| $\beta^A$ -globin minimal origin ΔCAAT+ ΔTATA #3 | 0.65 | 1.38 | 0.47 |
| Med14 pG4#4+5 origin #1 | 0.51 | 1.08 | 0.47 |
| Med14 pG4#4+5 origin #2 | 0.52 | 0.99 | 0.53 |
| Med14 pG4#4+5 minimal origin #1 | 0.52 | 1.15 | 0.46 |
| Med14 pG4#4+5 minimal origin #2 | 0.56 | 1.14 | 0.49 |

**Table 10: Transgene copy number determination in clonal cell lines.**
The table shows the qPCR results obtained with genomic DNA extracted from the clones selected for the experiments. For each clone, 2 ng of genomic DNA was amplified with a primer set amplifying a sequence within the construct (With) and another primer set amplifying a sequence 5 kb downstream from the insertion site for both alleles (Both). The ratio of the amounts of DNA obtained with the With and Both primer sets was used to determine transgene copy number in all clonal cell lines.
